# Integrative metabolome-genome analysis reveals the genetic architecture of metabolic diversity in sorghum grain

**DOI:** 10.1101/2025.10.14.682411

**Authors:** Deepti Nigam, Sarah Metwally, Songyue Shi, Priscilla Kolagani, Nasir Ali Khan, Ran Tian, Adil Khan, Melinda Yerka, Fang Chen, Yinping Jiao

**Author notes:** For correspondence: Yinping Jiao.

## Abstract

Natural variation in the grain metabolome plays a central role in shaping nutritional quality and end-use traits in grass crops. Understanding the genetic basis of this metabolic diversity is therefore essential, yet population-scale integration of metabolomics and genomics remains limited in sorghum, a climate-resilient C4 crop renowned for its exceptional heat and drought tolerance. Here, we integrated large-scale untargeted metabolomic profiling, population genomics, and artificial intelligence (AI)-based machine learning to systematically dissect grain metabolic diversity and its genetic architecture in sorghum. Untargeted metabolomic profiling of mature grains of the Sorghum Association Panel (SAP) identified 4,877 compounds, revealing extensive quantitative variation relevant to grain nutritional improvement. Metabolite-based genome-wide association studies (mGWAS) identified ∼4.15 million significant SNP–metabolite associations, revealing the heterogeneous genetic architecture of metabolic traits. Associated variants were enriched in genic and regulatory regions but depleted in intergenic regions, consistent with functional constraint. A total of 38 metabolite gene clusters revealed coordinated genetic control of core metabolic pathways. We further applied machine learning to identify key metabolites that underlie grain color variation and to prioritize associated candidate genes, demonstrating the utility of predictive models integrating genotype, metabolome, and end trait. Collectively, this work establishes a population-scale atlas of sorghum grain metabolomic and genetic diversity, available through the Sorghum Grain Metabolite Diversity Atlas (SorGMDA). This resource enables integrated metabolomics and genomic analyses and supports systems-level breeding strategies for improving grain nutritional quality.

## Introduction

Sorghum (*Sorghum* bicolor (L.) Moench) is a staple C4 monocotyledon crop in the *Poaceae* family (Upadhyaya et al. 2014). It exhibits high heat and drought tolerance, making it a cornerstone of agricultural systems in arid and semi-arid regions worldwide (Paterson et al. 2009; Ali et al. 2025). This crop is cultivated for human food, animal feed, and bioenergy production, but its full nutritional potential remains underutilized amid escalating global food demands (Liaqat et al. 2024). Sorghum is a gluten-free carbohydrate source rich in bioactive metabolites, including phenolic acids, flavonoids, and tannins (Tanwar et al. 2023). These molecules contribute to its high antioxidant capacity and potential health benefits, including anti-inflammatory (Burdette et al. 2010; Zhang et al. 2019) and anti-carcinogenic properties (Collins et al. 2024; Mazumder et al. 2024). These nutritional attributes uniquely position sorghum as a low-input crop that can be leveraged to address global food security and dietary health challenges, particularly in gluten-sensitive populations (Kaur et al. 2024).

Sorghum populations exhibit extensive genetic and phenotypic diversity (Boatwright et al. 2022), including substantial variation in grain bioactive metabolite concentrations (Kaufman et al. 2013; Habyarimana et al. 2019). Recent advances in sorghum breeding underscore the importance of grain bioactive metabolites (Boyles et al. 2017; Girard and Awika 2018; Vanamala et al. 2018; Przybylska-Balcerek et al. 2019), which are increasingly recognized for their dual roles in promoting human health (Farrar et al. 2008) and enhancing crop resilience and nutritional quality (Mukherjee et al. 2024). For instance, phenolic metabolites not only support human health by reducing the risk of chronic diseases, including diabetes, obesity, cancer, and cardiovascular disease (Awika et al. 2009; Dykes et al. 2013; Arbex et al. 2018; Girard and Awika 2018; Chen et al. 2021b), but also play key roles in plant defense against biotic and abiotic stresses by detoxifying reactive oxygen species (ROS) (Cingoz and Gurel 2016).

Elucidating the genetic basis of metabolite variation is critical for improving sorghum grain quality through molecular breeding and genetic engineering (Zheng et al. 2011). Genome-wide association studies (GWAS) have emerged as a powerful tool for dissecting the genetic architecture of grain quality traits in sorghum, including starch composition (Boyles et al. 2017; Chen et al. 2019), protein content (Li et al. 2018), and micronutrient diversity (Thakur et al. 2024). However, the resolution of these studies is often constrained by several factors, including the use of complex, integrative traits that may not capture underlying biochemical intermediates, the polygenic nature of many grain quality phenotypes (Hamblin et al. 2007), and extensive linkage disequilibrium (LD) in elite temperate sorghum germplasm around loci for height, maturity, cytoplasmic male sterility systems (the most common basis of heterotic groups), and photoperiod insensitivity. For example, starch content, a key determinant of grain quality (Thitisaksakul et al. 2012), is typically measured as a single trait, yet its biosynthesis involves intricate biochemical pathways regulated by multiple gene networks, including those controlling amylose synthesis and starch branching enzymes (López-González et al. 2019; Khan et al. 2024). Previous GWAS in sorghum grain have typically used composite traits such as total starch, protein, or oil content (Boyles et al. 2017; Rhodes et al. 2017). These traits represent composite biochemical phenotypes and therefore provide limited resolution for resolving their underlying genetic architecture (McCarthy et al. 2008; Rhodes et al. 2017).

The plant metabolome encompasses a diverse array of biochemical compounds, including starches, micronutrients, and bioactive secondary metabolites such as phenolic acids and flavonoids, which collectively determine its nutritional value, end-use functionality, and health benefits (Wurtzel and Kutchan 2016). Metabolomics-based genome-wide association studies (mGWAS) offer a powerful framework for linking metabolic traits to their genetic determinants by integrating high-resolution metabolite profiling with genome-wide genotyping (Luo 2015; Zhou et al. 2019; Chen et al. 2021a). As mGWAS relies on the same underlying genetic variants and statistical framework as conventional GWAS, it shares similar limitations (Tibbs Cortes et al. 2021), including challenges in identifying causal variants and potential confounding by genetic background and environmental factors (Tam et al. 2019; Merrick et al. 2022; Young 2024).

Recent advances in untargeted metabolome profiling utilizing high-performance liquid chromatography (HPLC) coupled with a Q Exactive Orbitrap mass spectrometer (Michalski et al. 2011) have revolutionized our ability to characterize complex metabolite mixtures with ultra-high resolution and mass accuracy (Eiler et al. 2017), enabling the effective separation and identification of structurally similar isomers (Siddique 2021; Qin et al. 2022). Unlike targeted metabolomics, which focuses on quantifying a predefined set of known metabolites, untargeted approaches aim to capture a broad, unbiased spectrum of metabolites, including metabolic intermediates and previously unknown or unexpected compounds, making them particularly valuable for exploratory studies and hypothesis generation (Ribbenstedt et al. 2018). LC–MS-based metabolomics studies in major cereal crops, including *Zea mays* (maize) (Baniasadi et al. 2014; Desmet et al. 2021), *Triticum aestivum* (wheat) (Ren et al. 2021; Hamade et al. 2024), and *Oryza sativa* (rice) (Xiao et al. 2018; Cheng et al. 2025), have been increasingly incorporated with chemometric and machine learning approaches, such as principal component analysis (PCA), partial least squares discriminant analysis (PLS-DA), and random forest classification, to support metabolite discovery, feature selection, and sample discrimination. Building on these analytical advances, metabolomics-based genome-wide association studies (mGWAS) have emerged as a powerful framework for linking metabolic variation to its underlying genetic determinants in cereal crops.

mGWAS has been widely applied to dissect metabolic traits across diverse cereal species. For instance, a multi-environment mGWAS of 983 metabolite features across 702 maize genotypes identified putative causal variants in five candidate genes linked to metabolic traits (Wen et al. 2014). In wheat, profiling of 805 metabolites across 182 accessions using >14,000 SNP markers revealed >1,000 significant associations and identified multiple candidate genes associated with variation in flavonoid metabolite substrates (Chen et al. 2020). In rice, mGWAS identified *Os02g57760,* a previously unknown nicotinic acid N-methyltransferase gene associated with grain size (Chen et al. 2016). Similarly, in barley, mGWAS identified associations between fructan composition and specific metabolic pathways and known fructan biosynthesis genes, further connecting metabolite profiles to grain quality traits (Matros et al. 2021). In foxtail millet, the integration of GWAS, mGWAS, and microbiome-wide association studies (MWAS) revealed genotype-dependent effects on root-associated microbial communities and metabolite profiles, which could help plant breeders select for desirable crop-microbiome interactions based on host genetics (Wang et al. 2022). Together, these advances underscore the potential of metabolome-assisted breeding to translate metabolic diversity into actionable targets for crop improvement (Fernie and Schauer 2009).

The Sorghum Association Panel (SAP) comprises genetically diverse accessions (Rhodes et al. 2017; Boatwright et al. 2022) with wide variation in grain quality traits (Boyles et al. 2017), making it an ideal resource for dissecting complex trait architecture. Recent deep whole-genome sequencing of the SAP has generated a high-density map of genetic variants, providing a robust foundation for high-resolution association mapping (Boatwright et al. 2022). Using this resource, we characterized the grain metabolome of 266 SAP accessions and explored the genetic basis of metabolite variation to develop a systems-level framework for metabolome-assisted breeding and nutritional improvement in sorghum.

## Results

### Grain metabolome profiling in the sorghum association panel (SAP)

To capture the metabolic diversity of sorghum grain, we performed untargeted metabolomic profiling on mature grains from 266 diverse accessions of the SAP. Metabolite annotation confidence was supported by high-quality MS/MS spectral matches (mass accuracy ≤ 5 ppm and spectral similarity score ≥ 0.7) against multiple databases, ensuring reliable metabolite identification for downstream genetic analyses (Supplemental Fig. S1). In total, 4,877 metabolites were detected, including 3,527 in positive-mode electrospray ionization (ESI+) and 1,350 in negative-mode electrospray ionization (ESI−). Among these, 89.6% were successfully annotated, 52.8% were assigned known biological functions, and 36.7% were mapped to established metabolic pathways (Supplemental Fig. S2A; Supplemental Table S1). Unless otherwise specified, metabolite identifications reported in this study are putative and correspond to Metabolomics Standards Initiative (MSI) levels 2–3. Accordingly, correlations among mass features are interpreted cautiously, as co-varying signals may reflect shared biochemical regulation or analytical artifacts rather than identical compound identities.

The quantitative reliability of the untargeted metabolomics dataset was validated using targeted LC–MS/MS quantification of 10 flavonoids and phenylpropanoid-derived compounds across 12 sorghum lines (36 samples; three biological replicates per line; Supplemental Fig. S2B; Supplemental Table S2). The analyzed metabolites included luteolinidin, apigeninidin, 7-methoxyapigenidin, luteolinidin-5-O-glucoside and apiforol, which contribute to sorghum pericarp pigmentation (Nip and Burns 1969; Wu et al. 2016), as well as additional flavonoids and phenylpropanoid derivatives such as naringenin, eriodictyol, luteolin, apigenin and 5,7-dihydroxycoumarin, which participate in broader defense-related and antioxidant pathways (Pontieri et al. 2021). Comparison of metabolite abundances obtained from targeted and untargeted approaches revealed strong concordance, with Pearson correlation coefficients ranging from 0.762 to 0.996 (Supplemental Fig. S2B; Supplemental Table S2), confirming the robustness of both relative and quantitative measurements.

Identified metabolites were classified into distinct biochemical categories, with carbohydrates (21%), benzenoids (17.5%), organoheterocyclic compounds (10.2%), fatty acyls (9.5%), and nucleic acids (9.3%) representing the five most abundant classes (Supplemental Fig. S3A). Comparative profiling across SAP accessions revealed substantial diversity in metabolite abundance patterns (Supplemental Fig. S3B), with subsets of accessions exhibiting similar global metabolite profiles. When grouped by biochemical class, metabolites such as amino acids, lipids, and polyphenols showed coordinated abundance patterns across accessions, reflecting class-specific variation within the metabolome (Supplemental Fig. S3C).

### Landscape of metabolite diversity in sorghum grain

From the 4,877 detected metabolites, 2,206 metabolites were retained for downstream analyses based on detection in >95% of the 266 SAP accessions and conformity to normality criteria (Shapiro–Wilk test, P > 0.05) (Supplemental Figs. S4A, B and S5A, B). The high-confidence metabolite set was used for all downstream analyses. Principal component analysis (PCA) revealed structured metabolomic variation across the SAP population, with the first two principal components explaining 42% and 38% of the total variance, respectively (Supplemental Fig. S5C). This high proportion of explained variance suggests coordinated variation among metabolites within shared biosynthetic pathways, consistent with biochemical and genetic co-regulation, rather than reduced metabolic diversity.

To characterize complementary aspects of metabolite variation across the SAP population, we evaluated the coefficient of variation (CV) as a measure of abundance dispersion, diversity indices (Shannon and Simpson) to capture metabolite richness and evenness (Buzas and Hayek 2005; Halliday and Rohr 2019), and repeatability (R) to assess measurement consistency across biological replicates. Following batch correction and median normalization (Supplemental Fig. S5D; Supplemental Table S3), CVs were calculated for 2,206 high-confidence metabolites. More than half of these metabolites (51.11%) exhibited high variability (CV > 80%), while 23.17% and 16.38% showed intermediate (CV = 60–80%) and moderate (CV = 40–60%) variability, respectively. Only 9.34% of metabolites displayed low variability (CV < 40%). The predominance of highly variable metabolites indicates substantial quantitative diversity within the SAP population, providing a strong foundation for mGWAS (Fig. 1A; Supplemental Table S1).

**Figure 1.**
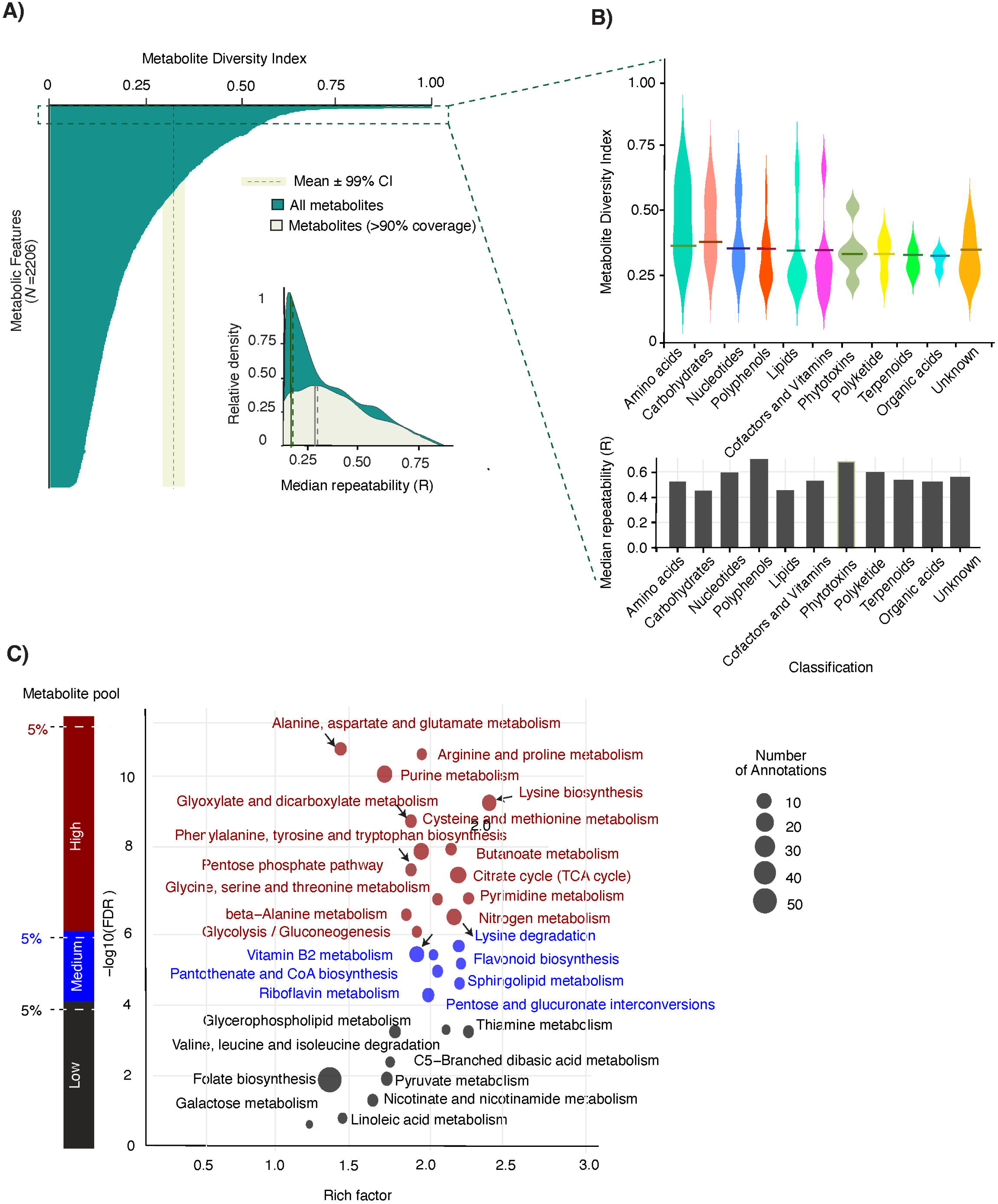
Sorghum grain metabolome diversity. (A) Distribution of the Shannon diversity index across metabolites, reflecting metabolite richness and evenness across samples. The Shannon diversity index is conceptually distinct from the coefficient of variation (CV), which measures dispersion. The inset shows the relative density distribution of median repeatability (R) for metabolites detected in >90% of samples. Dashed lines indicate the mean ±99% confidence interval. (B) Violin plots showing the distribution of the Shannon diversity index across metabolite superclasses. The bar plot below indicates median repeatability (R) within each superclass. (C) Bubble plot showing pathway enrichment among metabolite groups with high-, medium-, and low-diversity indices based on the Shannon diversity index. The x-axis represents enrichment magnitude (rich factor), defined as the ratio of significant metabolites mapped to a pathway relative to the total number of annotated metabolites in that pathway. The y-axis represents statistical significance [−log10(FDR)]. Bubble size indicates the number of annotated metabolites per pathway. Bubble colors denote pathways enriched in high-, medium-, and low-diversity metabolite groups. The side bar at left indicates the metabolite subsets used for enrichment analysis, corresponding to the top 5% high-diversity, middle 5% medium-diversity, and bottom 5% low-diversity metabolites

To focus on robust and consistently detected metabolites, diversity and repeatability patterns were further examined among metabolites detected in >95% of genotypes (coverage threshold). Consistent with these patterns, metabolites with greater abundance variability also exhibited higher heterogeneity in abundance distribution across genotypes, reflected by elevated diversity indices. The density distributions further indicated that highly prevalent metabolites largely showed moderate-to-high diversity together with strong repeatability (R), supporting the robustness and stability of metabolite quantification across genotypes, as illustrated by the repeatability density distribution and narrow confidence intervals in the inset plot (Fig. 1A). Broad-sense heritability (H²) estimates for individual metabolites further revealed substantial genetic contributions to metabolite variation across the population (Supplemental Table S3). In addition, the 99% confidence interval (CI) around diversity estimates further supported the robustness of the observed metabolite variation patterns across genotypes.

Metabolite superclasses, including amino acids, carbohydrates, nucleotides, polyphenols, polyketides, terpenoids, and organic acids, exhibited distinct distributions of diversity indices (Fig. 1B). Differences in median repeatability (R) were also observed across superclasses, indicating variation in measurement consistency. Together, these patterns suggest class-specific differences in metabolic diversity and underlying genetic control (Fig. 1B). Metabolites were further classified into high-, medium-, and low-diversity index groups across genotypes (Supplemental Table S2).

Pathway enrichment analysis showed that metabolites with high diversity index (top 5%) were predominantly associated with amino acid and central carbon metabolic pathways, including alanine, aspartate and glutamate metabolism, arginine and proline metabolism, purine metabolism, lysine biosynthesis, cysteine and methionine metabolism, and the citrate cycle (TCA cycle), which exhibited the strongest enrichment significance (−log10[FDR] > 8–10; Fig. 1C). Additional enriched pathways included glyoxylate and dicarboxylate metabolism, phenylalanine, tyrosine and tryptophan biosynthesis, pentose phosphate pathway, glycine, serine and threonine metabolism, β-alanine metabolism, glycolysis/gluconeogenesis, pyrimidine metabolism, and nitrogen metabolism. Metabolites within the medium-diversity group were enriched in pathways including vitamin B2 metabolism, flavonoid biosynthesis, pantothenate and CoA biosynthesis, sphingolipid metabolism, riboflavin metabolism, and pentose and glucuronate interconversions, generally exhibiting moderate enrichment significance (−log10[FDR] ≈ 4–6). In contrast, metabolites within the low-diversity group were associated with pathways such as glycerophospholipid metabolism, valine, leucine and isoleucine degradation, folate biosynthesis, pyruvate metabolism, nicotinate and nicotinamide metabolism, galactose metabolism, and linoleic acid metabolism, which displayed comparatively weaker enrichment significance (−log10[FDR] < 3–4) (Fig. 1C).

### Network architecture of the sorghum grain metabolome

Metabolites exhibiting correlated accumulation patterns are often co-regulated within shared biosynthetic pathways, suggesting coordinated metabolic control (Sawada et al. 2009; Fukushima et al. 2011). To identify co-regulated molecular clusters, we constructed correlation-based networks using the 2,206 high-confidence metabolites (Fig. 2).

**Figure 2.**
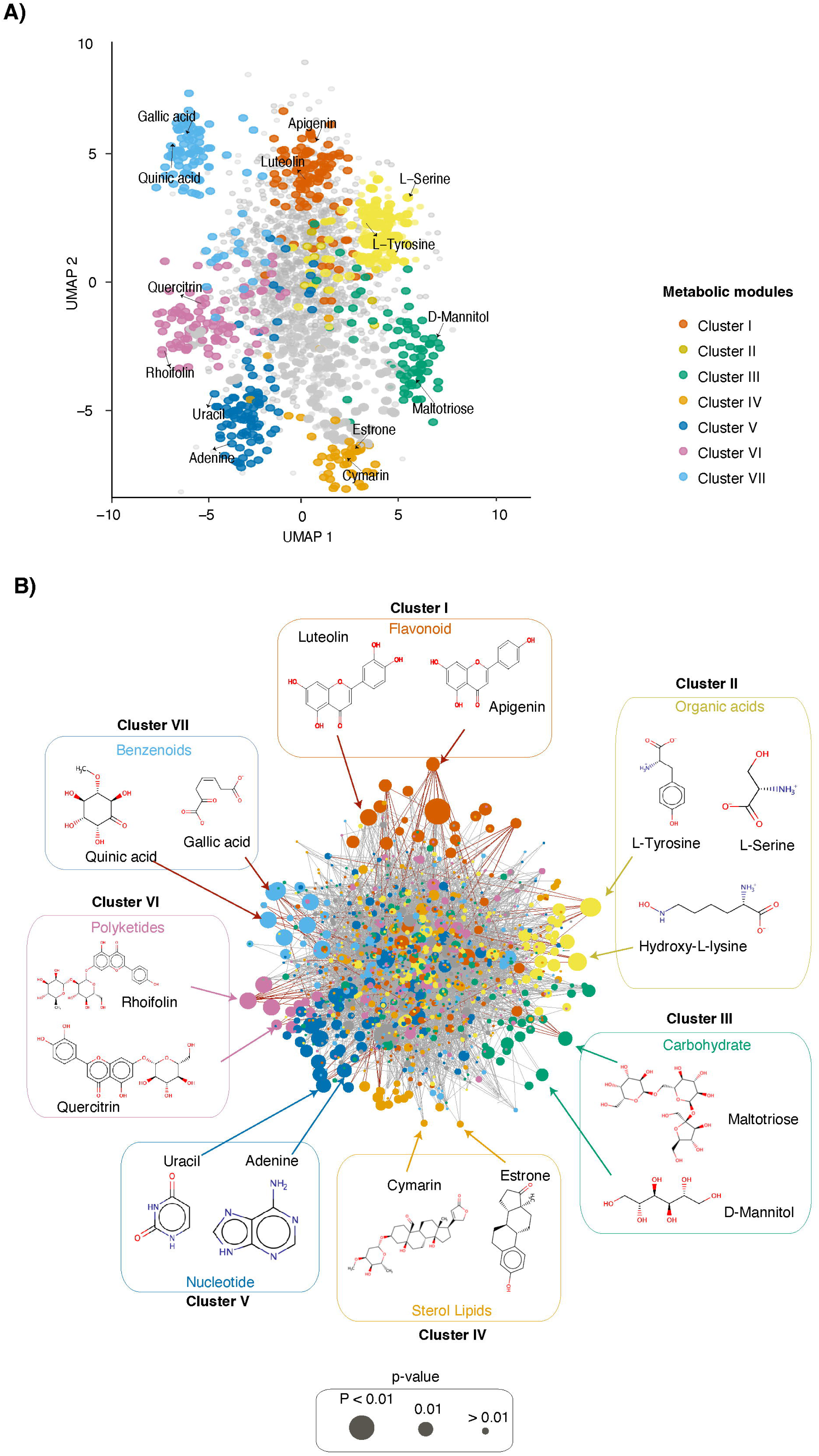
Sorghum grain metabolomic network. (A) UMAP visualization of 2,206 sorghum grain metabolites colored according to the seven metabolite modules identified in the network. Labeled metabolites indicate representative compounds within each module. (B) Metabolite association network constructed from pairwise molecular MS/MS similarity. Each node represents a metabolite signal. Node size reflects p-values from the Kolmogorov–Smirnov test: larger nodes (p < 0.01) indicate higher statistical significance, whereas smaller nodes (p > 0.01) indicate lower statistical significance. Gray edges indicate molecular MS/MS relationships based on spectral similarity, whereas red edges highlight co-accumulating grain metabolites with similar accumulation patterns across 266 sorghum accessions. These edge types represent distinct, non-overlapping relationships. Metabolite clusters are outlined, and representative metabolite structures are shown for each cluster.

To examine the global organization of metabolite associations, we generated a Uniform Manifold Approximation and Projection (UMAP)-based low-dimensional representation using metabolite module membership and network topology (Fig. 2A). The UMAP projection revealed partially overlapping yet distinct neighborhoods corresponding to seven metabolite modules, indicating that coordinated metabolite variation is organized into structured biological communities rather than arising from random associations.

Consistent with the UMAP organization, the metabolite association network resolved distinct clusters associated with both primary metabolism (e.g., carbohydrates, organic acids, and nucleotides) and specialized metabolism (e.g., flavonoids, benzenoids, and polyketides). The network was constructed by linking signals (nodes) with similar accumulation profiles at a significance threshold of *P* ≤ 0.01 (Fig. 2B). In the network, red edges indicate co-accumulating metabolites, whereas grey edges represent structural similarity inferred from MS/MS data. Functional annotation of these modules revealed strong biochemical coherence among clustered metabolites.

For instance, Cluster I (flavonoids) contained metabolites such as luteolin (peak ID F121951, m/z = 286.0477) and apigenin (peak ID F6485, m/z = 270.05), whose identities were supported by reference MS/MS spectra and characteristic flavone fragmentation patterns (Supplemental Table S1). Additional clusters were similarly characterized by shared MS/MS spectral features and metabolite class enrichment. Notably, metabolites such as L-serine, L-tyrosine, and hydroxy-L-lysine co-clustered within a single organic acid-related module (Fig. 2B). Although these compounds differ structurally and functionally, their co-clustering may reflect shared structural features that contribute to similar fragmentation behavior, coordinated regulation within amino acid metabolism, or participation in related biochemical processes. Alternatively, their grouping may reflect coordinated regulation within amino acid metabolism or participation in related biochemical processes, highlighting the ability of network-based metabolomics to uncover previously unrecognized biochemical relationships.

Although most modules displayed strong biochemical coherence, some highly correlated mass features carried apparently unrelated putative annotations. Such relationships are common in large-scale untargeted metabolomics datasets and may arise from shared MS/MS fragmentation patterns, co-eluting compounds, or limitations in database-based annotation, particularly for metabolites assigned at MSI confidence levels 2–3 (Domingo-Almenara et al. 2017; Sindelar and Patti 2020). Consequently, network correlations primarily represent evidence of coordinated metabolite variation rather than definitive structural similarity. For example, highly correlated features assigned to distinct chemical classes or apparently exogenous compounds may reflect common accumulation patterns, shared regulatory mechanisms, or annotation ambiguity rather than direct biochemical relationships (Chaleckis et al. 2019; Zhou et al. 2022).

### Genetic variation associated with metabolome diversity in sorghum grain

To explore the genetic architecture underlying metabolic diversity, mGWAS was performed using 38.8 million high-quality SNPs (minor allele frequency > 0.05) obtained from whole-genome resequencing of the SAP panel (Boatwright et al. 2022).

As positive controls, GWAS signals were recovered for known genes associated with two representative metabolites, (**-**)-epigallocatechin (C□□H□□O□; putatively identified, MSI level 2) and D-(**-**)-fructose (C□H□□O□; putatively identified, MSI level 2) demonstrating the robustness of our metabolome profiling and mGWAS framework (Supplemental Fig. S6, Supplemental Table S4). (-)-Epigallocatechin (EGC) is a flavan-3-ol that functions as an antioxidant in plants and has been implicated in defense-related responses, reactive oxygen species (ROS) scavenging, cell-wall strengthening, and inhibition of fungal growth in various plant species (Grace 2005). Although direct functional studies in sorghum remain limited, its role as a precursor of condensed tannins supports a potential contribution to grain protective mechanisms (Bröhan et al. 2011; Rao et al. 2018; Wu et al. 2019).

The mGWAS of EGC identified 47 significant SNPs (*P* ≤ 10^-8^), including genetic variants located ∼14 kb downstream (*P* = 4.85 × 10□□□), of the well-characterized *Tannin1* (*Tan1*, *Sobic.004G280800*) gene on Chromosome 4 (Supplemental Fig. 6A and Supplemental Table S4). *Tan1* encodes a WD40-repeat transcription factor that regulates the expression of genes in the proanthocyanidin (condensed tannin) biosynthetic pathway, seed coat pigmentation, and pathogen defense (Wu et al. 2012; Wu et al. 2019). An additional signal was detected ∼15 kb upstream of *TRANSPARENT TESTA 6* (*TT6*, *Sobic.003G418000*) on Chromosome 3 (*P* = 1.60 × 10□¹□), which encodes flavanone 3-hydroxylase (F3H), a 2-oxoglutarate-dependent Fe(II) oxygenase (Supplemental Fig.6A and Supplemental Table S4). F3H catalyzes the stereospecific hydroxylation of flavanones to dihydroflavonols, an essential step in flavonoid biosynthesis, as observed in *Arabidopsis thaliana* (Pelletier and Shirley 1996), rice (Lam et al. 2014) and maize (Falcone Ferreyra et al. 2015). As a second positive control, D-fructose, a simple sugar central to primary metabolism and energy storage, is a major component of the sorghum grain carbohydrate profile and yield (Almodares and Hadi 2009; Simeone et al. 2017). Our mGWAS identified 29 significant SNPs (P ≤ 10□□), including a major association peak on chromosome 4 located approximately 28 kb upstream of Starch Synthase IIb (*SSIIb*; *Sobic.004G238600*, *P* = 1.53 × 10□¹□, Supplemental Fig. S6B), a well-characterized enzyme involved in starch biosynthesis (Campbell et al. 2016).

Using the 2,206 high-confidence metabolites, mGWAS identified 4,151,451 unique SNP associations at a significance threshold of *P* ≤ 1 × 10□□, of which 2,221,701 passed robust processing based on a resample model inclusion probability (RMIP) ≥ 0.05 (Supplemental Figs. S7A, B; Supplemental Table S5). In this resampling-based framework, SNPs with higher RMIP values were considered more reliable for association mapping (Bian and Holland 2017).

To further prioritize robust associations, we quantified the number of significant SNPs per metabolite feature using varying genome-wide significance thresholds and RMIP cutoffs (Supplemental Fig. S7B). The distribution is strongly right skewed, with a median of 7 SNPs per metabolite and 90% of metabolites associated with ≤17 SNPs. In contrast, a small subset of metabolites accounts for disproportionately large numbers of associations, with the top 1% exceeding 45,000 SNPs (Supplemental Fig. S7C). These results indicate that SNP–metabolite associations are unevenly distributed and driven in part by highly connected metabolites, with potential implications for downstream enrichment analyses.

Among the SNPs identified through mGWAS, many were located near known biosynthetic enzymes, while others represented novel associations, highlighting the utility of mGWAS for uncovering previously unrecognized genetic mechanisms. For example, lysine ranks among the top 5% most diverse metabolites within the SAP (Fig. 1C). The mGWAS of lysine (C□H□□N□O□; putatively identified, MSI level 2) revealed 77 significant SNPs (*P* < 10□□) (Fig.3A, Supplemental Table S4), including four loci located within ±50 kb of genes encoding key enzymes in the lysine biosynthetic pathway. Accordingly, Supplemental Table S4 reports candidate genes located within this ±50 kb window rather than all genome-wide association signals. These included three aspartokinases (AK) (*Sobic.001G011700*, *P* = 6.65 × 10□¹², *Sobic.003G412100*, *P* = 1.93 × 10□□ and *Sobic.002G155100*, *P* = 4.49 × 10□□), dihydrodipicolinate synthase (DHDPS, *Sobic.006G186600*, *P* = 3.00 × 10□□), and lysine decarboxylase (LDC, *Sobic.001G127700, P* = 3.93 × 10□□), each corresponding to distinct steps in the pathway (Fig.3B). Three of these enzymes revealed specific allelic variants at their respective loci: a CC-to-TT transition on Chromosome 3 for AKIII, TT-to-GG on Chromosome 6 for DHDPS, and CC-to-GG on Chromosome 1 for LDC (Fig.3C).

**Figure 3.**
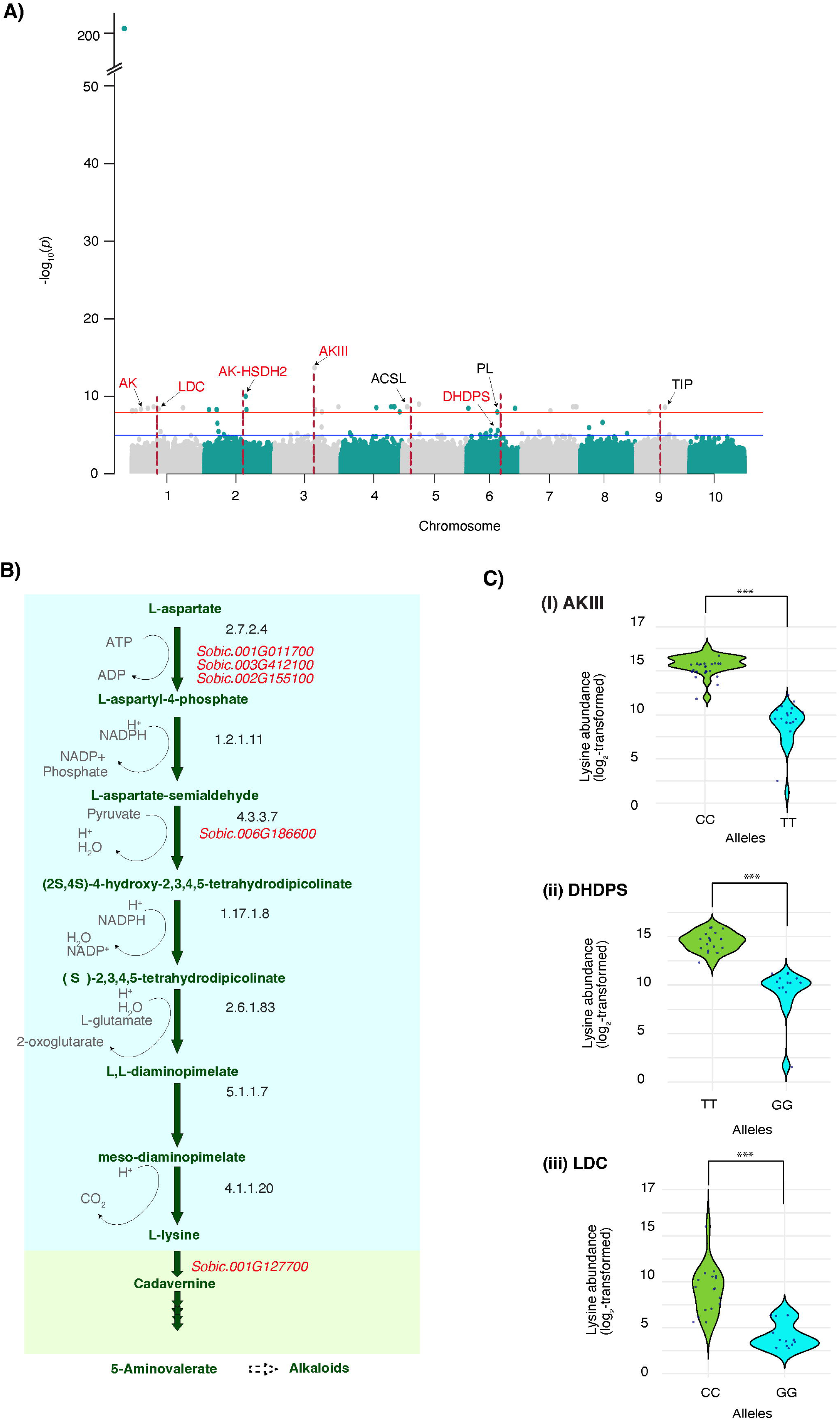
Genetic dissection of lysine accumulation in sorghum grain. (A) Manhattan plot showing metabolite-based genome-wide association study (mGWAS) results for lysine (C□H□□N□O□) accumulation. The x-axis represents the 10 sorghum chromosomes, and the y-axis represents statistical significance [−log10(P)]. Associations were assigned to genes located within ±50 kb of lead SNPs. Red dashed lines indicate loci near candidate genes, including AK (aspartate kinase) and LDC (lysine decarboxylase) on chromosome 1; AK-HSD2 (aspartate kinase–homoserine dehydrogenase 2) on chromosome 2; AKIII (aspartate kinase III) on chromosome 3; ACSL (acyl-CoA synthetase long-chain family member) on chromosome 5; DHDPS (dihydrodipicolinate synthase) and PL (pectate lyase-related gene) on chromosome 6; and TIP (tonoplast intrinsic protein) on chromosome 9. (B) Lysine biosynthetic pathway in sorghum showing major intermediates, enzymatic steps, and corresponding genes. Enzyme Commission (EC) numbers are provided where applicable. Genes corresponding to key mGWAS candidate loci are highlighted in red. (C) Lysine accumulation comparison of the alleles of three candidate genes: AKIII, DHDPS, and LDC. The y-axis represents log2-transformed lysine abundance, and each point represents an individual sorghum accession grouped by allele.

These allelic variants were associated with significant differences in lysine accumulation, indicating that allelic variation at these loci contributes to the natural variation in lysine content observed across the SAP. AK catalyzes the first committed step from L-aspartate, DHDPS catalyzes a key branch-point reaction in the diaminopimelate pathway, and LDC converts L-lysine to cadaverine, linking primary lysine metabolism to alkaloid biosynthesis (Zhu and Galili 2003; Jander and Joshi 2009; Galili et al. 2016; Bunsupa et al. 2017). In addition to the known genes, mGWAS also identified genetic variants located in regions distant from any previously characterized gene involved in lysine biosynthesis. For example, we identified a SNP ∼40 kb upstream of PECTATE LYASE 1-RELATED (*PL-1*; *Sobic.006G014400*) on Chromosome 6 (*P* = 3.00 × 10□□) (Fig.3A, Supplemental Table S4). *PL-1* is known for its role in cell wall modification and fruit softening (Xu et al. 2022), suggesting a potential indirect link to lysine accumulation through modulation of cell wall-associated metabolic processes.

### Genomic distribution of Metabolite-Associated Variants

To identify genomic regions enriched for metabolite-associated variation, we compared SNPs from two datasets: the input set of genome-wide SNPs in the SAP panel and the significant mGWAS SNPs. Comparison between the input SNPs (n = 43,983,694) and mGWAS SNPs (n = 4,151,451) revealed shifts in the genomic distribution of associated variants (Fig. 4A, B; Supplemental Table S6). Notably, mGWAS SNPs were relatively enriched in genic and regulatory regions, including 5′ untranslated regions (5′-UTR) and promoter-proximal upstream regions. Fold-enrichment analysis indicated significant overrepresentation of missense (8.6-fold; *P* = 6.32 × 10□²□), synonymous (7.9-fold; *P* = 8.61 × 10□¹□), and UTR variants (∼2-fold; 5′-UTR: *P* = 7.27 × 10□□; 3′-UTR: *P* = 1.07 × 10□¹□) (Fig. 4C; Supplemental Table S6). Moderate enrichment was observed for upstream, downstream, and intronic variants (< 2-fold), whereas intergenic variants were modestly depleted (0.78-fold; log□ enrichment ≈ −0.37), indicating that not all genomic categories were enriched.

**Figure 4.**
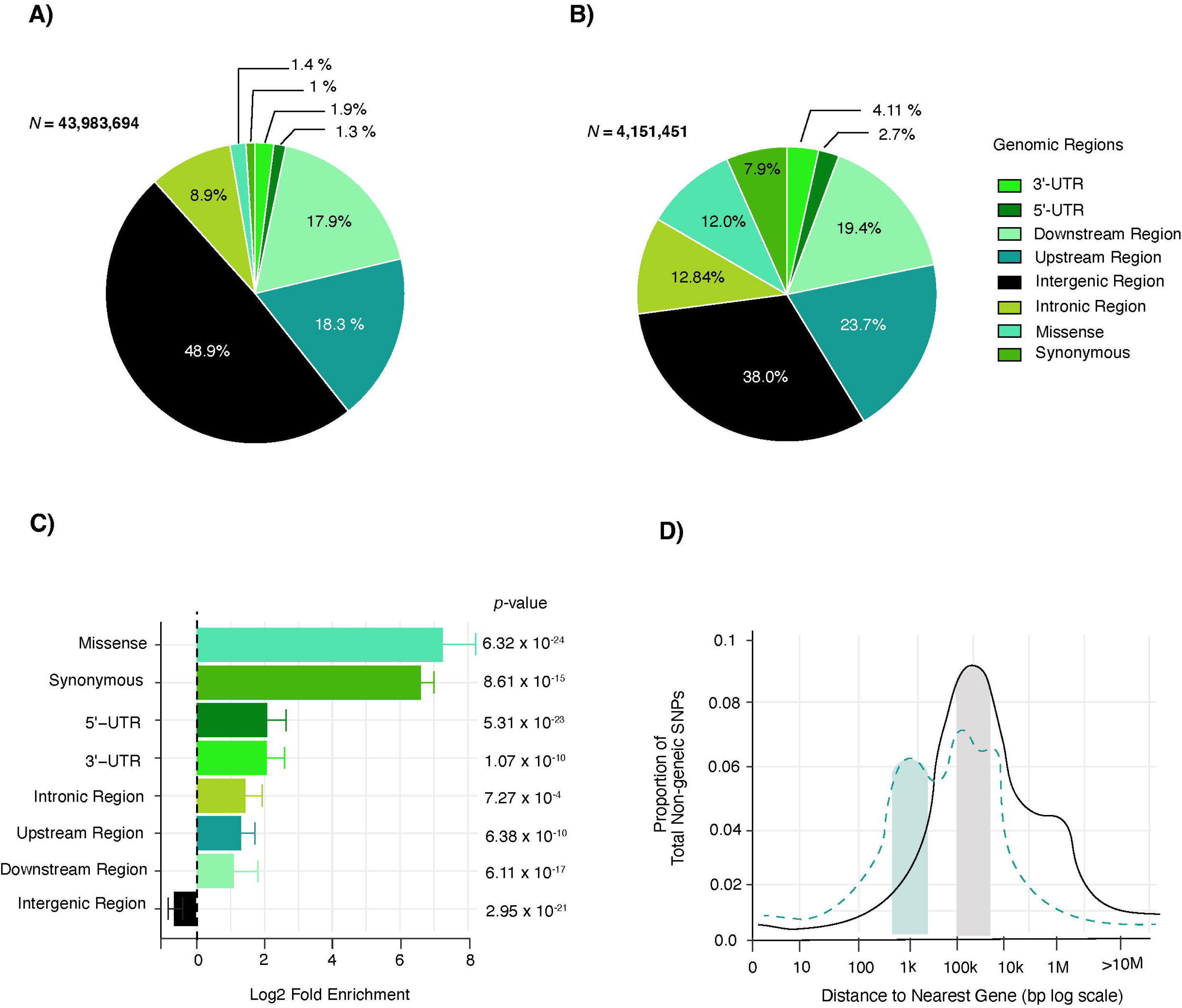
Genomic enrichment of significant SNPs identified through mGWAS. (A) Distribution of SNPs across genomic regions in the SAP input dataset. (B) Distribution of SNPs significantly associated with 2,206 metabolites identified through mGWAS. Percentages in (A) and (B) do not sum to 100% because functional annotations (e.g., missense and synonymous variants) overlap with genomic location categories and are therefore not mutually exclusive. (C) Enrichment of SNPs across genomic regions, calculated as log□(mGWAS proportion / input proportion). Positive values indicate enrichment, whereas negative values indicate depletion relative to the background input SNP dataset. (D) Distance distribution of non-genic SNPs to the nearest gene (log scale). The solid line represents the observed distribution of mGWAS-associated SNPs, and the dashed line represents the background distribution from the input dataset. Shaded regions highlight enrichment patterns, with the green region indicating SNPs proximal to genes and the gray region indicating more distal SNPs.

To investigate the spatial distribution of non-genic metabolite-associated variants, the distance from each significant non-genic mGWAS SNP to its nearest annotated gene was quantified (Fig. 4D; Supplemental Table S6). Significant non-genic SNPs showed enrichment near genes relative to the genomic background, with a major peak within ∼1 kb of the nearest gene, consistent with promoter-proximal regulatory regions. A secondary peak was observed at ∼100 kb, which may reflect distal regulatory elements (Fig. 4D. In contrast, SNPs located far from genes (>1 Mb) were depleted relative to background non-genic variants. Together, these patterns indicate that metabolite-associated variants preferentially localize to genic and nearby regulatory regions, supporting a model in which metabolic diversity is primarily shaped by functionally relevant loci rather than by distant non-coding regions with limited evidence of regulatory activity.

### Metabolic gene cluster (MGC) variants in sorghum grain

Given the enrichment of metabolite-associated variants within gene-dense and regulatory regions, we next examined whether these variants preferentially localize to higher-order genomic features involved in metabolic regulation, particularly MGCs. MGCs comprise physically co-located genes that encode enzymes functioning in the same, co-regulated metabolic pathway, and they play key roles in determining the nutritional and biochemical composition of sorghum grains (Nützmann et al. 2016; Liang et al. 2021). Across the sorghum genome, we identified 38 MGCs distributed across 10 chromosomes, collectively harboring 434 biosynthetic genes (Fig.5A and Supplemental Fig. S8A; Supplemental Table S7). These clusters belong to diverse metabolic classes, including polyketide, lignin, lignan-polyketide, lignin-saccharide, lignin-terpene, amino acids, saccharide-polyketide, saccharide-terpene, alkaloid, and terpene. Notably, these clusters were located within intergenic distances ranging from 26.95 kb to 806.04 kb, and Chromosome 5 harbors the highest number of gene clusters (eight in total), including five polyphenol, one amino acid, and one carbohydrate metabolic cluster, suggesting that this chromosome region serves as a hub for secondary metabolic pathways (Fig.5A).

**Figure 5.**
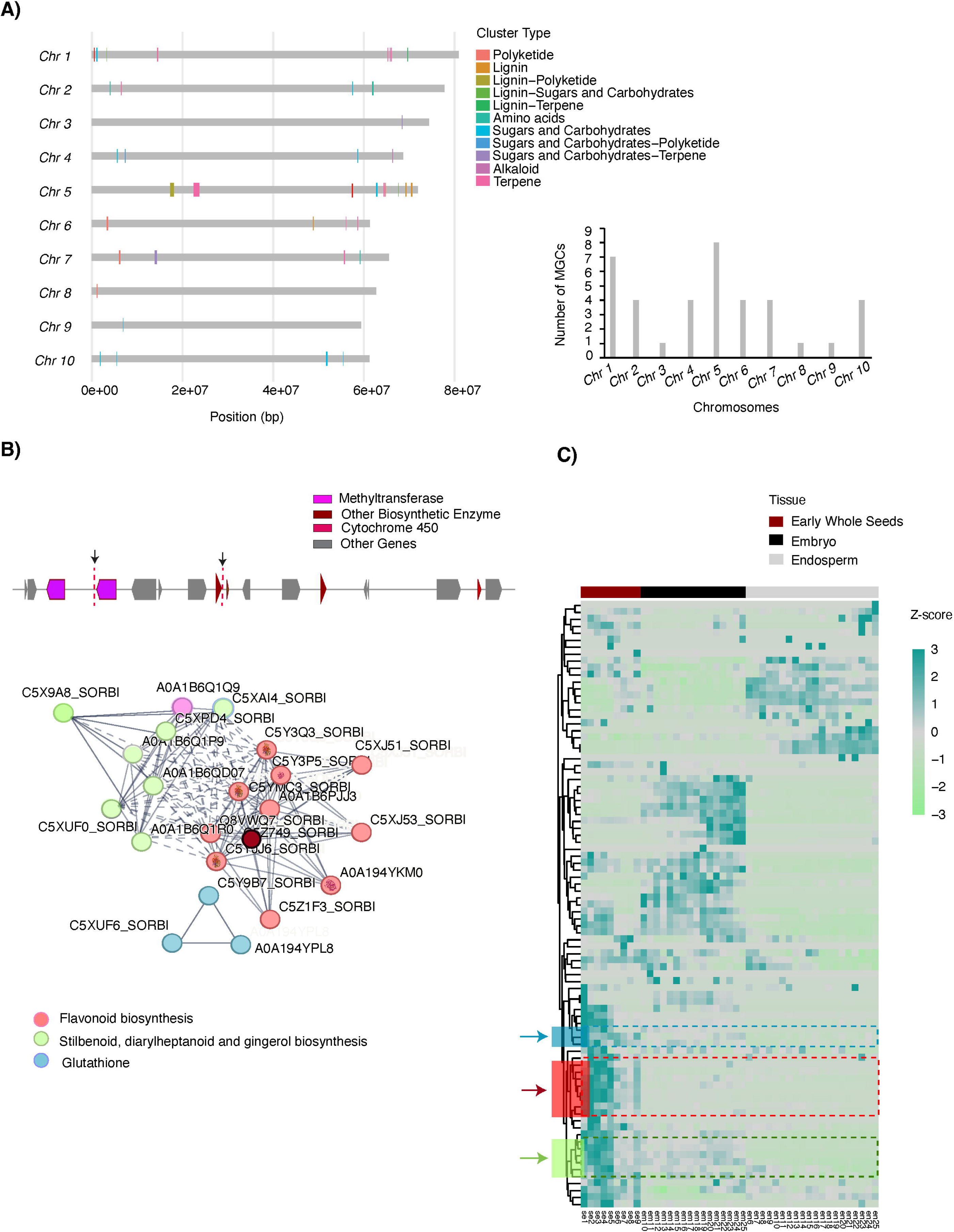
Chromosomal distribution and functional analysis of metabolic gene clusters (MGCs). (A) Distribution of MGCs across 10 sorghum chromosomes. Cluster are color-coded by biosynthetic pathway. The x-axis represents chromosomal position (base pairs), and the y-axis indicates chromosome number. The inset bar plot shows the total number of MGCs identified per chromosome. (B) The lignin metabolic gene cluster. The top panel depicts the gene arrangement and location of sequence variants near methyltransferase-encoding genes and a dirigent enzyme gene. The bottom panel shows the co-expression network of genes within this cluster, highlighting functionally related genes involved in flavonoid biosynthesis (red nodes), stilbenoid, diarylheptanoid, and gingerol biosynthesis (green nodes), and glutathione biosynthesis (blue nodes). A dark red central node (C5Z749_SORBI) serves as a major hub within the flavonoid biosynthesis subnetwork and is extensively connected to multiple surrounding pink flavonoid-related nodes, indicating strong co-expression relationships among these genes. (C) Heatmap showing the normalized expression profiles of co-expressed lignin biosynthesis genes (FPKM > 1). Genes involved in flavonoid biosynthesis are marked in blue, while genes associated with stilbenoid, diarylheptanoid, and gingerol biosynthesis are highlighted in red and green, respectively.

To evaluate the distribution of mGWAS signals across MGCs, we quantified SNP–cluster associations across multiple genomic windows. In total, 135, 173, and 249 SNP–cluster associations were identified within 50 kb, 100 kb, and 500 kb windows, respectively, reflecting the expected increase in associations with broader genomic intervals (Supplemental Fig. S8B). Enrichment analysis, expressed as log2(observed/expected), indicated that metabolite-associated SNPs are generally overrepresented within MGC regions relative to genome-wide expectations, although the degree of enrichment varied substantially among clusters (Supplemental Fig. S8B). This pattern supports a non-random but heterogeneous localization of mGWAS signals across metabolic gene clusters.

To prioritize likely causal variants and minimize indirect associations, we focused on 85 high-confidence SNPs located within ±50 kb of biosynthetic genes (Supplemental Table S7). These SNPs were distributed across 35 of the 38 MGCs, with several localized in clusters associated with phenylpropanoid and lignin metabolism, including Cluster 17 (lignin–polyketide), Cluster 24 and Cluster 23 (lignin), Cluster 22 (lignin-sugars and carbohydrates), and Cluster 19 (sugars and carbohydrates metabolism). These clusters harbor key biosynthetic domains such as dirigent proteins, cytochrome P450s, chalcone synthases, and methyltransferases, which are known to play central roles in lignin and flavonoid biosynthesis. These prioritized SNPs and clusters were subsequently used for downstream functional and co-expression analyses (Fig. 5B and C; Supplemental Table S7).

We next examined SNPs associated with variation in lignin content within MGCs. Lignin plays a critical role in seed development (Frei 2013) and contributes to tolerance against abiotic and biotic stresses (Li et al. 2022), including insect herbivory and pathogen infection (Dowd et al. 2016). A total of 20 genes located within ±50 kb of the lignin-associated variants, including methyltransferase loci and a nucleoporin-related gene, exhibited seed-specific expression and strong co-expression with other genes involved in specialized metabolism, collectively forming a distinct functional network (Fig.5B). Within this network, two major metabolic modules were identified: one related to flavonoid biosynthesis (7 genes) and another encompassing stilbenoid, diarylheptanoid, and gingerol biosynthesis (10 genes), representing a broader phenylpropanoid-derived pathway (Fig. 5B). Within the flavonoid module, C5Z749_SORBI (*Sobic.010G210700*), annotated with anthocyanidin reductase activity (GO:0033729), exhibited the highest connectivity and served as a central hub linking multiple flavonoid-associated genes within the co-expression network. (Fig. 5B; Supplemental Table S8). A third, smaller module comprising three genes associated with glutathione metabolism was also detected; however, this module showed no correlation with the other two (Fig.5B). Expression profiling during seed development revealed three distinct gene clusters, indicating coordinated and tissue-specific expression patterns (Fig. 5C; Supplemental Table S7 and Supplemental Table S8). The enrichment of specialized metabolic pathway genes within this co-expression network, together with their coordinated expression patterns, suggests that genetic variation near methyltransferase loci may contribute to broader regulatory control of specialized metabolism during seed development.

### Metabolic regulation of sorghum grain color

Metabolomic diversity underlies phenotypic variation in complex end-use traits and enables detailed dissection of their biochemical bases (Carreno-Quintero et al. 2013; Karakas et al. 2025; Rai et al. 2025). Here, we use grain color as a representative trait to investigate the metabolic regulation of this key seed trait in sorghum. Grain color in sorghum and other cereals is a complex trait determined by the morphology of the pericarp and endosperm tissues, as well as by the accumulation of diverse classes of pigment-related metabolites, including flavonoids (e.g., anthocyanins), carotenoids, and phlobaphenes (Dykes et al. 2009; Su XiaoYu et al. 2017; Kathuria et al. 2024). Given the complex, nonlinear nature of grain color variation, we applied machine learning models (Tiozon et al. 2023; Zhang et al. 2023b) to identify relationships between metabolite profiles and grain color classes recorded in the GRIN database (Volk and Richards 2008). This approach complements mGWAS by linking metabolite variation to observable grain color phenotypes.

Metabolomic profiles from 211 SAP accessions were analyzed across four grain color categories: brown (77 lines), red (22 lines), yellow (10 lines), and white (102 lines) (Supplemental Table S9). Hierarchical clustering of metabolite abundances across grain color groups revealed distinct metabolic signatures, including differences in the composition of 49 secondary metabolites (Fig. 6A, Supplemental Fig. S9A, Supplemental Table S10). These metabolites were predominantly polyphenols and exhibited clear accumulation patterns corresponding to grain color variation. PCA further supported a key role for polyphenols in driving grain pigment diversity, with PC1 and PC2 explaining 43% and 37% of the variance among accessions, respectively (Supplemental Fig. S9B). Feature selection analyses identified seven top-ranked polyphenols (all putatively identified; MSI levels 2–3) that showed high feature importance and thus, were strongly associated with grain pigmentation: 3-coumaric acid, 6-hydroxy-7-methoxycoumarin, genistein, isorhamnetin-3-glucoside, isorhamnetin-3-rutinoside, kaempferol, and quercetin-3-(6’’-malonyl)-glucoside (Fig. 6B; Table S1; Supplemental Table S10; Supplemental Fig. S9C). These metabolites belong primarily to the phenolic acid, coumarin, isoflavone, and flavonol classes. Notably, none of the top-ranked metabolites belonged to the flavan-3-ol or proanthocyanidin (condensed tannin) subclasses, which are the major polyphenol groups associated with bitterness and astringency in sorghum (Awika and Rooney 2004; Kobue□Lekalake et al. 2007; Wu et al. 2012).

**Figure 6.**
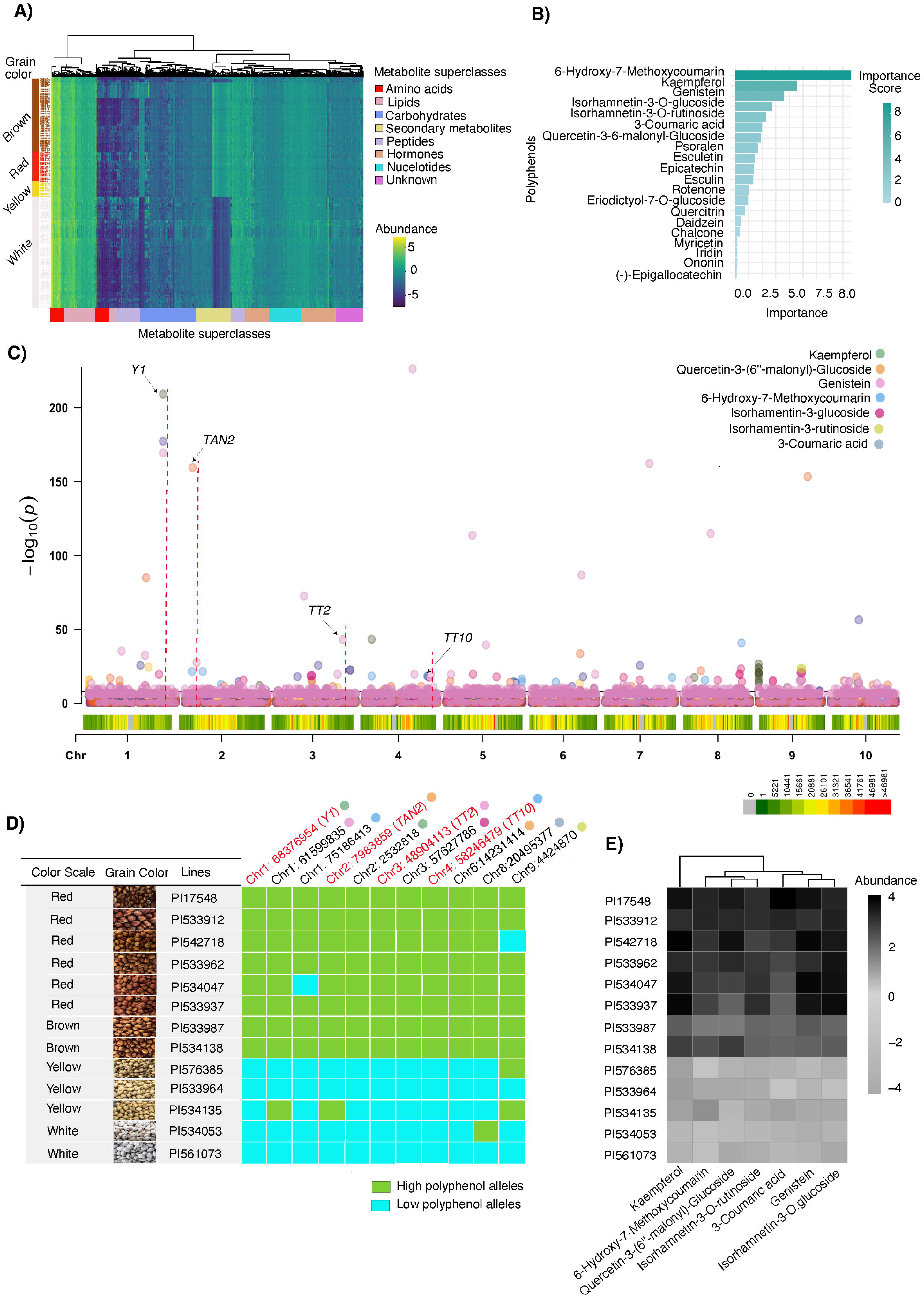
Metabolomic clustering and genomic loci associated with sorghum grain color variation. **(A)** Hierarchical clustering of 2,206 metabolites profiled across 211 sorghum accessions grouped according to grain color: brown (77 accessions), red (22 accessions), yellow (10 accessions), and white (102 accessions). Metabolite superclasses are color-coded to highlight distinct metabolic signatures among grain color groups. **(B)** Feature importance plot showing the top-ranked polyphenols identified by Random Forest (RF) and XGBoost models based on predictive importance scores. **(C)** Composite Manhattan plot of the seven top-ranked polyphenols showing significant genomic loci associated with metabolite variation. The color gradient represents genome-wide SNP density estimated using consecutive genomic windows in CMplot (bin size = 1 Mb), with increasing color intensity corresponding to higher SNP density across genomic regions. Previously reported genes were annotated at their corresponding chromosomal positions. **(D)** Distribution of genotypes for the 11 significantly associated SNPs linked to polyphenol variation across sorghum accessions with distinct grain colors. Green blocks indicate alleles associated with high polyphenol accumulation, whereas blue blocks indicate alleles associated with low polyphenol accumulation. **(E)** Normalized abundance of seven key polyphenols across randomly selected sorghum lines, illustrating their combined genetic effects on grain pigmentation.

ROC curve analysis evaluated the discriminatory power of these metabolites for grain color classification (Supplemental Fig. S10; Supplemental Table S11). Among them, isorhamnetin-3-glucoside (Bordiga et al. 2014; Sytar et al. 2019; Coklar and Akbulut 2021) and genistein (Bursać et al. 2017; Lee et al. 2017) showed the strongest statistical separation among the four grain color groups, consistent with their roles as flavonoid-derived pigments within the phenylpropanoid pathway that directly influence grain pigmentation. Moderate classification performance was observed for 3-coumaric acid (Noel et al. 2005), 6-hydroxy-7-methoxycoumarin (Kai et al. 2006), and quercetin-3-(6’’-malonyl)-glucoside (Ojwang et al. 2012), which are phenylpropanoid and flavonol intermediates that may contribute more broadly to secondary metabolism rather than specifically to grain color contrast. In contrast, kaempferol (Park et al. 2006) showed limited discriminatory potential, indicating that its abundance is either relatively similar across color classes or that it plays a more general antioxidant or structural role not tightly coupled to visible grain pigmentation. Alternatively, kaempferol may be distributed across multiple grain tissues rather than being confined to the seed coat, thereby weakening its association with grain color (Goufo and Trindade 2014; Mbanjo et al. 2020; Mbanjo et al. 2023).

The mGWAS of these seven polyphenols identified 224 significant genetic variants (*P* < 10□¹□) associated with grain color variation (Fig. 6C; Supplemental Table S12). A total of 214 genes were mapped within a 50-kb flanking region of significant mGWAS signals, including genes previously implicated in pigment regulation, such as *Y1* (*Sobic.001G398100*, *P =* 7.76 × 10^-210^), *Tan2* (*Sobic.002G076600*, *P =* 3.24× 10□^160^), *TT2* (*Sobic.003G183800*, *P =* 2.55× 10□^49^), and *TT10* (*Sobic.004G236000*, *P =* 3.57 × 10□^22^) as well as enzymes known to regulate the phenylpropanoid, flavonoid, and lignin biosynthetic pathways (Supplemental Fig. S11A, Supplemental Table S12). The QQ plot showed a clear deviation from the expected null distribution, reflecting an enrichment of significant associations across all seven polyphenols analyzed (Supplemental Fig. S11B). This pattern underscores the robustness of the mGWAS approach in elucidating the polygenic control underlying variation in grain color. The proximity of these genes to mGWAS signals suggests that genetic variation in regulatory or coding regions may modulate polyphenol accumulation, thereby influencing differential pigmentation among sorghum accessions.

From the joint analysis of the 11 significant SNPs identified from mGWAS of the seven top-ranked polyphenols across sorghum grain color groups (Table 1), we found dark-colored grains (red and brown) consistently exhibited higher accumulation of polyphenols compared with light-colored grains (yellow and white). The combination of polyphenol-associated alleles linked to higher accumulation was more frequent in red (71–78%) and brown (9.8–22%) sorghum accessions, whereas it occurred at lower frequencies in yellow (4.6–10%) and white (4–8.5%) accessions (Table 1, Supplemental Fig. S12; χ² = 13.82, df = 3, *P* = 0.00316). These results suggest that sorghum lines carrying higher polyphenol-associated alleles across the 11 SNPs tend to exhibit darker grain color.

**Table 1.** Allelic variation at candidate SNPs associated with polyphenol accumulation in sorghum grains. Notes: *Violin plots represent the distribution of log□-transformed metabolite abundance between allele groups for each SNP–metabolite association. Asterisks indicate statistically significant differences between allele groups (\**P* < 0.05, \*\**P* < 0.01, \*\**P* < 0.001).

To further evaluate the combined effects of these loci and metabolites on grain color variation, we analyzed genotype patterns and metabolite abundance profiles in 13 randomly selected sorghum accessions representing a gradient from white to dark red grains. The genotype distribution across the 11 significant SNPs showed clear differentiation between dark- and light-colored accessions, with alleles associated with higher polyphenol accumulation enriched in darker grains (Fig. 6D). Consistently, normalized abundance profiles of the seven key polyphenols revealed elevated metabolite accumulation in dark-colored accessions relative to light-colored accessions (Fig. 6E). Together, these findings demonstrate a strong association among SNP allele combinations, polyphenol accumulation, and grain pigmentation, highlighting the utility of integrated metabolomic and genomic analyses in dissecting complex agronomic traits.

## Discussion

### Genetic architecture of the sorghum grain metabolome revealed by mGWAS

This study revealed remarkable chemical diversity of sorghum grain in SAP, with more than 4,800 detected features, including 2,206 high-confidence metabolites that covered 95% of the SAP population across 266 accessions (Supplemental Fig. S4). By integrating untargeted metabolomics with mGWAS, we systematically characterized metabolite–SNP associations and identified causal candidate genes for several nutritionally and agronomically important pathways.

In maize, metabolome-scale GWAS identified 1,459 locus–trait associations across 983 grain metabolite features (Wen et al. 2014). In rice, metabolome-based GWAS dissected the genetic basis of the rice metabolome and identified hundreds of metabolite associations (Gong et al. 2013). In wheat, metabolite-based GWAS of 805 grain metabolites detected 1,098 associations and nominated candidate genes for flavonoid decoration (Chen et al. 2020). In barley, GWAS on metabolite accumulation in wild barley NAM populations identified mQTLs associated with sugar metabolism (Gemmer et al. 2021). Collectively, these studies illustrate the scale and resolution achievable with metabolome-enabled genetic mapping and provide a comparative framework for interpreting our results. Within this context, the analysis of sorghum grain color as a complex endpoint phenotype further demonstrates how integrating metabolomic and genetic data with machine learning–based predictive modeling can resolve the molecular basis of agronomic traits.

The strength of mGWAS lies in its ability to move beyond traditional phenotypic selection toward direct selection on biochemical composition (Song et al. 2022). In sorghum, this is particularly relevant for traits that are difficult or expensive to phenotype (e.g., antioxidant polyphenols, free lysine, condensed tannins) yet critically influence nutritional quality, food functionality, and pre-harvest sprouting resistance. Comparable successes in other crops- including maize vitamin E content (Lipka et al. 2013), rice 2-acetyl-1-pyrroline (2-AP) aroma (Li et al. 2024), tomato lycopene (Sauvage et al. 2014), and wheat phenolic acids (Chen et al. 2020)-illustrate that metabolite-guided breeding can rapidly deliver consumer- and climate-oriented varieties.

To facilitate data accessibility and application, we developed SorGMDA, a public interactive atlas linking sorghum grain metabolite profiles, mGWAS results, and candidate genes. This resource enables breeders and geneticists to query metabolites of interest, identify accessions with extreme concentrations of beneficial compounds, and retrieve linked genetic variants. For example, (–)-epigallocatechin can be explored in SorGMDA to identify high-accumulating lines and their associated candidate genes for breeding purposes (Supplemental Fig. S13). Increasing grain (–)-epigallocatechin content is valuable due to its antioxidant activity, contributions to seed defense, and potential benefits for nutritional quality and human health, making it an important target for sorghum improvement (Awika and Rooney 2004; Mokra et al. 2022). This resource can facilitate the translation from metabolomic and genomic data into practical breeding tools.

Despite these advances, challenges remain. Metabolite levels are highly sensitive to the environment, and our study was conducted on field-grown material from a single season and location, so future multi-environment trials will be essential to dissect G×E interactions and develop robust, environment-resilient markers. Second, although untargeted metabolomics provides extraordinary coverage, absolute quantification and full structural elucidation of all features remain resource intensive. In addition, metabolite annotations assigned at MSI levels 2–3 based on database matching represent putative identities and should be interpreted cautiously, particularly when evaluating correlations among highly associated mass features, as covariance in signal intensity does not necessarily indicate identical structural identity.

### Functional interpretation of mGWAS signals and candidate genes

Our mGWAS identified more than four million genetic variants associated with metabolite traits, including both known loci and previously uncharacterized regions. Similar observations in other crop mGWAS studies have highlighted the importance of regulatory variation and gene networks in controlling metabolic traits (Wen et al. 2014; Chen et al. 2020; Matros et al. 2021).

The identification of known loci such as *Tannin1, TT6, and SSIIb* demonstrates the robustness and high quality of both the metabolomic and genetic datasets. Notably, significant variants were enriched in promoter-proximal and 5’-untranslated regions, highlighting the prominent role of cis-regulatory variation in shaping metabolite abundance. In addition, the identification of metabolic gene clusters containing co-expressed biosynthetic enzymes underscores the importance of physical clustering in the coordinated regulation of specialized metabolic pathways. The large number of uncharacterized candidate genes uncovered by this study provides a valuable opportunity to expand our understanding of the biosynthetic networks governing sorghum grain composition. However, the functional characterization of these genes remains a long-term challenge requiring systematic, multifaceted investigation. To support community-wide efforts, we have made our mGWAS results openly available through the SorGMDA database, providing a foundation for downstream functional analyses.

Dissecting gene function will require integrating complementary approaches. Fine mapping of associated loci can refine candidate gene lists and identify causal variants. Subsequent multi-layered analyses, including transcriptomics, metabolomics, proteomics, and targeted biochemical assays, can elucidate the mechanistic roles of candidate genes within metabolic pathways. Ultimately, functional validation through genetic manipulation, such as gene knockouts or overexpression, will be essential to confirm causal relationships and assess their potential utility in crop improvement. For example, in cotton, integrative metabolomic and transcriptomic analyses identified two MYB transcription factors as key regulators of 252 metabolites associated with fiber development and other agronomic traits (Zhang et al. 2025). Similarly, in tomato, metabolome-enabled GWAS combined with transcriptomic, and population genetic analyses identified five major loci controlling fruit flavor. Subsequent breeding efforts targeting these loci demonstrated the power of metabolome-assisted breeding for improving complex quality traits (Zhu et al. 2018). Collectively, these post-GWAS studies will advance our understanding of sorghum specialized metabolism and establish a framework for translating genetic discoveries into precision breeding and metabolic engineering strategies to improve grain quality and nutritional value.

## Methods

### Sample collection

The original SAP was obtained from the U.S. National Plant Germplasm System (NPGS) via the Germplasm Resources Information Network (GRIN). Plants were grown at the Texas Tech University Research Farm in Lubbock, Texas (33°35′52.9″N, 101°54′21.4″W; elevation 992 m). The site is characterized by a semi-arid climate, with an average annual rainfall of approximately 469 mm, and fine-loamy, mixed, superactive, thermic Aridic Paleustalf soils typical of the Amarillo series. Supplemental irrigation (∼2.56 cm per week) was provided throughout the growing season. Plants were grown under field conditions and were randomized within the field to minimize potential spatial effects. Panicles were covered with pollination bags prior to flowering to prevent outcrossing. For metabolite profiling, a subset of 266 SAP accessions was randomly selected based on their ability to reach physiological maturity within the local growing season. For each accession, we prepared three biological replicates, with 20 seeds in each replicate. Each sample was freeze-dried for 12 h using a benchtop freeze dryer (Labconco FreeZone 2.5 Liter, -84 °C) connected to a vacuum pump (Labconco Model 117) to prevent degradation. To minimize heat generation during milling, tubes were frozen in liquid nitrogen before grinding. The dried grains were then finely ground in 2-mL microcentrifuge tubes using a bead-mill homogenizer.

### Untargeted UHPLC-MS metabolite profiling and data processing

For metabolite extraction, 35 mg of each freeze-dried sample was transferred into 2-mL microcentrifuge tubes containing two ceramic zirconium oxide beads. Samples were immersed in a pre-cooled extraction solvent consisting of methyl tert-butyl ether (MTBE) and methanol (3:1, v/v), supplemented with internal standards: 100 μL of 1,2-diheptadecanoyl-sn-glycero-3-phosphocholine (for lipid analysis) and 50 μL each of corticosterone and ampicillin (for polar metabolite analysis). Samples were homogenized using a Precellys 24 bead mill at 5000 Hz for two 45-s intervals. Homogenates were incubated on an orbital rotator at 100 rpm for 45 minutes at room temperature to facilitate extraction. Phase separation was induced by adding a water/methanol mixture (3:1, v/v), followed by additional homogenization and centrifugation at 20,000 × *g* for 5 minutes at 4 °C. The resulting upper (lipid-containing) and lower (polar metabolite) phases were transferred into separate tubes, dried using a SpeedVac, or stored at -80 °C for future analysis. Prior to UHPLC-MS analysis, the lipid extracts were reconstituted in acetonitrile:2-propanol:water (65:30:5, v/v), and the polar extracts in UPLC-grade methanol:water (1:1, v/v). Both fractions were centrifuged to remove particulates and transferred into autosampler vials. To monitor instrument stability and analytical repeatability, a mixed quality control (QC) sample was prepared by pooling 20 μL aliquots from all samples across accessions and biological replicates.

Metabolite separation and detection were performed in both ESI positive and negative ion modes using reversed-phase liquid chromatography (RPLC) on a Vanquish UHPLC system (Thermo Fisher Scientific) equipped with a Waters Acquity UPLC HSS T3 column (1.8 µm, 2.1 × 100 mm), coupled to a Q Exactive HF high-resolution mass spectrometer (Thermo Fisher Scientific, USA) for metabolic profiling. The mobile phases consisted of 0.1% formic acid in water (A) and 0.1% formic acid in methanol (B), delivered at a flow rate of 0.40 mL/min. The gradient program was as follows: 0.5% B initially; 0.5–50% B from 5.5–6 min; 50–98% B from 6–12 min; held at 98% B from 12–13 min; and re-equilibrated at 0.5% B from 13–15 min. The injection volume was 5 µL.

The resulting raw mass spectrometry (MS) data (.raw) files were converted to .mzXML format using MSConvert, with an MS-level filter applied (Chambers et al. 2012). Metabolite quantification was achieved by analyzing signal intensities using the XCMS and CAMERA data processing pipelines (Smith et al. 2006). A minimum sample threshold of three was set during the XCMS grouping phase to capture rare metabolites across this diverse sample set. Missing values were handled using median imputation and interpolation methods to preserve data integrity and ensure completeness. Mass features identified by XCMS-CAMERA were filtered based on retention times (60-630 seconds) and precise mass (*m/z* < 0.5 at the first decimal point), excluding naturally occurring isotopes. Although false annotations were frequent, MS adducts were retained to minimize the risk of discarding true metabolites. Feature quantification was adjusted for tissue fresh weight and normalized against the total ion count of each sample to account for technical variability. The resulting data were log-transformed to enable multivariate analyses. To further refine the dataset, interquartile range filtering and Pareto scaling were applied before conducting statistical analysis. All additional statistical assessments and visualizations were performed using the ggplot2 package in R (Kassambara 2013). The metabolic trait (m-trait) data for the SAP were calculated as the average of three biological replicates measured using UPLC-MS. Specifically, for each metabolite m (where *m*=1,2,…,4,877) in each accession l (where *l*=1,2,…,266), the m-trait value *Mm,l* was computed as:

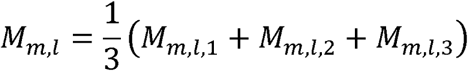

where *Mm,l*,1, *Mm,l*,2, *Mm,l*,3, are the normalized metabolite intensities from the three biological replicates.

### Metabolite identification confidence and validation

Metabolite annotation was conducted using multiple complementary databases, including HMDB (Wishart et al. 2022), MassBank (Horai et al. 2010), ChemSpider (Pence and Williams 2010), and KEGG (Kanehisa and Goto 2000), enabling broad metabolite assignment across diverse chemical classes. Due to the large-scale, population-level nature of this untargeted metabolomics study and the lack of a comprehensive set of authentic standards for sorghum grain metabolites, no MSI Level 1 identifications were performed. Instead, all metabolites were assigned putative identities corresponding to Metabolomics Standards Initiative (MSI) Levels 2-3 (Sansone et al. 2007; Fiehn et al. 2008; Spicer et al. 2017). MSI Level 2 assignments were supported by high-resolution MS/MS spectral similarity to reference databases, while MSI Level 3 annotations were based on precursor mass accuracy and diagnostic fragmentation patterns consistent with specific metabolite classes. Confidence in biologically important metabolites, such as proanthocyanidin monomers, flavonoids, and phenolic acids-was strengthened through multiple lines of evidence, including high MS/MS spectral similarity scores (> 0.85), high-resolution precursor mass accuracy (≤ 5 ppm), co-localization of mGWAS signals with previously characterized biosynthetic loci (e.g., *Tannin1*), and agreement with published metabolome datasets. Together, these criteria provide robust and reproducible metabolite annotation suitable for downstream genetic and biochemical analyses while maintaining broad, unbiased coverage of the sorghum grain metabolome.

To assess biological reproducibility, correlations among the three biological replicates were evaluated prior to downstream analyses. Samples with replicate correlation coefficients below 0.90 were excluded from further analysis. For retained accessions, metabolite abundance values from the three biological replicates were averaged (mean) to generate a single representative value for each metabolite per accession for subsequent statistical and genetic analyses. The high reproducibility observed across retained biological replicates indicates that metabolite variation was primarily driven by genetic differences among accessions rather than micro-environmental or field-related variation.

### Targeted quantification of flavonoids using parallel reaction monitoring

To quantify a group of flavonoids in sorghum seeds, polar extracts from 12 selected lines, each analyzed in three biological replicates, were subjected to targeted Parallel Reaction Monitoring (PRM) analysis using a Thermo Scientific Vanquish UHPLC system coupled to a Q Exactive HF high-resolution mass spectrometer (Thermo Fisher Scientific, Waltham, MA, USA). Chromatographic separation was performed using a reversed-phase liquid chromatography (RPLC) method on a Waters Acquity UPLC HSS T3 column (1.8 µm, 2.1 × 100 mm). The mobile phases consisted of 0.1% formic acid in water (A) and 0.1% formic acid in methanol (B), delivered at a flow rate of 0.40 mL/min. The gradient program was as follows: 5% B initially; 5–50% B from 1–12 min; 50–98% B from 12–13 min; held at 98% B from 13–16 min; returned to 0.5% B from 16–17 min; and re-equilibrated at 0.5% B from 17–20 min. The injection volume was 5 µL. PRM data were acquired in both positive and negative ionization modes. Instrument parameters included a resolution of 30,000, AGC target of 3 × 10□, maximum injection time of 100 ms, and an isolation window of 1.0 m/z. Compound identification and quantification were based on accurate mass, retention time, and compound-specific precursor–product ion transitions. Method performance was assessed for calibration linearity, and relative quantification was performed using external calibration curves. Quality control samples were analyzed throughout the analytical sequence to monitor signal stability and instrument performance, ensuring data reliability across sample batches.

### Phenotypic data normalization and heritability estimation

The 2,206 distinct metabolite traits were normalized using the bestNormalize R package (v1.4.3), which systematically evaluates a range of transformation techniques, including Lambert W × F, Box-Cox, Yeo-Johnson, and Ordered Quantile, to identify the method that best improves normality, as measured by the Pearson *P* test statistic. Given the dataset’s high dimensionality, manual outlier curation was infeasible. Instead, a rules-based automated approach was employed to remove extreme values. After transformation, values exceeding ±1.5 times the interquartile range (IQR) from the first or third quartile of each trait distribution were considered outliers and treated as missing. Broad-sense heritability (*H²*) was then estimated for each metabolite using replicate-level averages per genotype. Each genotype was represented by three biological replicates, and values beyond predefined biological thresholds were excluded prior to heritability estimation. A linear mixed model was fitted, treating genotype as a random effect, as follows:

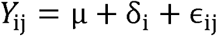

where *Y*_*ij*_ represents the normalized metabolite value for the *j*-th replicate of the *i*-th genotype, μ is the overall mean, *δ*_*i*_ is the random effect associated with the *i*-th genotype, and *ϵ*_*ij*_ is the residual error term. Variance components (*σ*_*G*_^*2*^ for genotype and *σ*_*R*_^*2*^ for residual error) were extracted from the model, and broad-sense heritability was calculated as:

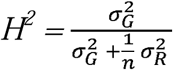

where n is the number of biological replicates per genotype (here, n= 3).

Repeatability (R) (Shurubor et al. 2005; Shurubor et al. 2007) was also calculated using the same variance components to assess the consistency of metabolite measurements across biological replicates, defined as:

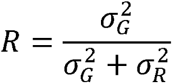

### Metabolite diversity analysis

Following normalization, metabolite intensities were scaled to ensure comparability across samples. Metabolites detected in fewer than 5% of the samples were excluded to reduce noise. For each sample, the relative abundance of each metabolite (*p*_*i*_) was calculated by dividing its intensity by the total intensity of all metabolites in that sample. To assess metabolic diversity, we employed two complementary indices: the Shannon Diversity Index and the Simpson Diversity Index (DeJong 1975; Gorelick 2006). Metabolite diversity was quantified in two ways: (a) metabolite richness, defined as the total number of metabolites detected in each accession; and (b) abundance-weighted diversity, which accounts for both the number and relative abundances of metabolites across accessions. These indices were selected to capture different aspects of diversity. The Shannon method (Pallister et al. 2017; Menni et al. 2020) was used to emphasize richness and evenness across metabolites, while the Simpson method highlighted dominance (Bi et al. 2021; Salgado et al. 2023). The Shannon Diversity Index (*H’*) was computed as:

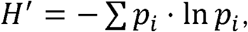

where pi represents the relative abundance of metabolite i, with higher *H’* values indicating greater diversity.

The Simpson Diversity Index (D) was calculated using the formula:

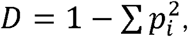

where *pi* represents the proportional abundance of the ith metabolite in an accession. This index gives more weight to abundant metabolites. Higher D values indicate that a few metabolites dominate the sample, while lower values suggest a more even distribution across metabolites. The index was computed for each accession, and the distribution of D values across all accessions was visualized using density plots.

### UMAP visualization and metabolite association network

To investigate the global organization of sorghum grain metabolites, a complementary Uniform Manifold Approximation and Projection (UMAP) visualization and MS/MS-based molecular association network were generated. The UMAP visualization was generated in R using the uwot package (Melville 2019) to represent metabolite module organization in a low-dimensional space based on network topology and module membership relationships. UMAP was used as an exploratory visualization approach to illustrate relative proximity and organization among metabolite classes and was not used for primary module identification. The metabolite association network was generated using MS/MS spectral similarity and metabolite co-accumulation relationships across 266 sorghum accessions.

Molecular networking analysis was performed using the Global Natural Products Social Molecular Networking (GNPS) platform (Aron et al. 2020). Nodes represent metabolite features, whereas edges indicate either molecular spectral similarity derived from MS/MS fragmentation patterns or correlated metabolite accumulation profiles. Metabolite modules were identified based on network topology and spectral relationships. Network visualization and annotation were performed in Cytoscape v3.10.1(Shannon et al. 2003). Node size was scaled according to statistical significance derived from the Kolmogorov–Smirnov test, with larger nodes indicating higher statistical significance (*p* < 0.01).

### Metabolite-based genome-wide association study (mGWAS)

Only the 38,873,876 SNPs from the 266 SAP accessions after filtering minor allele frequencies (≥ 0.05) were used for mGWAS (Boatwright et al. 2022). Metabolites were selected for GWAS if they were detected in at least 90% of the 266 phenotyped accessions and passed normality criteria. GWAS was independently performed on each of the 2,206 qualifying metabolites using a mixed linear model (MLM) (Zhang et al. 2010) with principal components (PCs) and the kinship matrix and one multi-locus model, Fixed and random model Circulating Probability Unification (FarmCPU) (Liu et al. 2016). These models were implemented with the R packages GAPIT (Lipka et al. 2012) and rMVP (v1.0.1) (Yin et al. 2021). In the MLM, a kinship matrix was integrated and implemented in rMVP. FarmCPU was run with the first three PCs as covariates, and the kinship matrix was calculated internally by the algorithm as a random effect. FarmCPU was executed with maxLoop = 10 and the MVP option for bin selection using method.bin = “FaST-LMM” (Lippert et al. 2011). The variance component analysis method, vc.method, was set to “GEMMA” (Team et al. 2024).

Bonferroni corrections were applied based on the effective number of independent markers in each dataset, calculated using the genetic type I error calculator (GEC v0.2) (Li et al. 2012). A stringent genome-wide significance threshold of 1×10□□ was used, corresponding to the Bonferroni-corrected cutoff to control the family-wise error rate. Due to instability in individual FarmCPU runs, the model was rerun 100 times with 90% subsampling and Resample Model Inclusion Probability (RMIP) was calculated for each variant. Only variants with RMIP > 0.05 were considered robust and retained for downstream interpretation. Manhattan plots were generated using the CMplot R package (v3.6.2), with SNP density estimated using the default 1-Mb genomic window size (bin.size = 1e6) (Yin et al. 2021).

To define independent association signals, a two-step procedure was applied. First, statistically significant SNPs located within 1 Mb on the same chromosome were merged into preliminary association peaks to capture broader genomic regions of association and reduce redundancy among closely spaced signals. Second, these peaks were refined into discrete loci using linkage disequilibrium (LD) information. Based on the observed genome-wide LD decay of ∼50 kb (r² = 0.2) in this population, significant SNPs within ±50 kb were grouped into single loci. Within each locus, lead SNPs were identified by accounting for LD among variants, ensuring that only independent association signals were retained.

Population structure and kinship were accounted for using principal components and kinship matrices, which reduced false associations, as confirmed by Q-Q plots (Voorman et al. 2011). Candidate genes within these defined loci were annotated using the BTx623 reference genome (v3.1) from Phytozome v14 (Goodstein et al. 2012), and genomic regions containing SNPs were further annotated using the Ensembl Variant Effect Predictor (VEP) (McLaren et al. 2016).

### Identification of metabolite gene clusters

Metabolic (biosynthetic) gene clusters (MGCs) in sorghum were identified using the plantiSMASH web server (version 2.0-beta4) according to default analysis protocols (Kautsar et al. 2017). Advanced parameters included a CD-HIT cutoff of 0.5 to cluster protein sequences with ≥50% identity and a minimum of 2 unique biosynthetic domains per candidate cluster. Built-in profile Hidden Markov Models (pHMMs) enhanced the detection of plant-specific biosynthetic domains. Candidate clusters were further manually validated based on their domain composition and gene co-expression patterns, using seed development RNAseq data (Khan et al. 2024). Significant mGWAS variants were then mapped onto the validated clusters to identify metabolite-associated loci and the genes harboring these variants.

### Expression profiling of lignin-associated genes during seed development

To demonstrate the potential of mGWAS to elucidate trait architecture through integration with time-resolved multi-omics data, we incorporated previously published RNA-seq expression profiles (E-MTAB-13406) (Khan et al. 2024) spanning sorghum seed development. The RNA-seq dataset was generated using the sorghum reference genotype BTx623 under field conditions at the same research farm.

This dataset provides high-resolution temporal expression profiling across 1–25 days post-anthesis (dpa) in multiple tissues, including embryo, endosperm, and whole seed. We leveraged the full temporal resolution of the RNA-seq data to characterize gene expression dynamics during seed development and to investigate lignin biosynthesis across tissues. Protein–protein interaction networks were inferred using STRING v12.0 (Von Mering et al. 2005), and correlation networks among genes were visualized using Cytoscape v3.10.1(Shannon et al. 2003).

### Machine learning-based analysis of grain color-associated metabolites

Grain color information for the SAP accessions was obtained from the U.S. National Plant Germplasm System (NPGS) (Council et al. 1990; Byrne et al. 2018). According to the Germplasm Resources Information Network (GRIN) database naming convention, grain color was categorized as follows: 1 = white, 2 = red, 3 = yellow, and 4 = brown, encompassing a total of 211 accessions. Prior to modeling, the metabolomic dataset (1,428 polyphenolic features) was pre-processed as follows: probabilistic quotient normalization, log□ transformation, removal of features with >20% missing values across samples, and k-nearest neighbor imputation (k = 5) for the remaining missing values using the impute R package.

To identify key polyphenolic metabolites associated with variation in seed color, we applied supervised machine learning models-Random Forest (RF) (Liaw and Wiener 2002; Erban et al. 2019) and Extreme Gradient Boosting (XGBoost) (Chen et al. 2017), in a multi-class classification framework (four color classes). These models used metabolite abundance data as input features and were trained to predict the grain color category. For full reproducibility, random seeds were fixed (np.random.seed(42), random.seed (42), and XGBoost seed = 42). This ensures that the ML methods are fully repeatable. Each metabolite was assigned an importance score by the models, indicating how strongly it contributes to distinguishing grain color categories. Metabolites with higher scores have a greater influence on the model’s classification performance. Models were trained and evaluated using repeated stratified 10-fold cross-validation (10 repeats, yielding 100 total folds) via RepeatedStratifiedKFold (scikit-learn), preserving the original class distribution in each fold (∼50% white, ∼25% red, ∼15% yellow, ∼10% brown) and mitigating class imbalance effects (Fontanari et al. 2022), where one class (e.g., positive or negative cases) is underrepresented relative to the other, which can bias model training and evaluation. Random Forest was run with the following hyperparameters: n_estimators = 2000, max_features = ‘sqrt’, min_samples_split = 2, min_samples_leaf = 1, bootstrap = True, class_weight = ‘balanced_subsample’. Feature importance was calculated using the mean decrease in Gini impurity averaged across all trees and all 100 folds. XGBoost was trained with objective = ‘multi:softprob’, num_class = 4, eta = 0.05, max_depth = 6, min_child_weight = 3, subsample = 0.8, colsample_bytree = 0.8, gamma = 0.1, alpha = 0.01, lambda = 1.0, tree_method = ‘hist’, and early_stopping_rounds = 50.

Feature importance was based on the “gain” metric, again averaged over the 100 folds. Metabolites that consistently ranked in the top 30 most important features in ≥70% of the cross-validation folds in both RF and XGBoost models were considered robust discriminators of grain color (Supplemental Table S7). Receiver Operating Characteristic (ROC) curve analysis (Hajian-Tilaki 2013; Burke 2023) was subsequently performed in R v4.3.1 using the pROC package to evaluate the discriminatory ability of each of the 27 individual polyphenolic metabolites across the four color groups (one-vs-rest and pairwise comparisons). The area under the curve (AUC) and 95% confidence intervals were calculated using 2000 bootstrap replicates.

Lead SNPs associated with seven polyphenolic compounds were then analyzed to assess allelic variation. Alleles were numerically encoded as follows: the reference allele was assigned a value of 1, the alternative allele a value of 2, and the heterozygous allele a value of 1.5. Haplotype analysis was performed using the R package geneHapR (Zhang et al. 2023a). Functionally linked SNPs were extracted and analyzed for their correlation with phenotypic traits using the SKAT package (Lee et al. 2011; Wu et al. 2011). Differential metabolomic abundance for each allele was visualized using the ggplot2 package (Wickham 2011). Joint haplotypes were constructed by clustering haplotype blocks spanning multiple genomic regions.

Subsequent association analyses examined the relationships between these joint haplotypes and candidate polyphenolic metabolites implicated in grain color variation, assessing their potential functional relevance. Additionally, a genome-wide correlation network analysis was conducted to identify SNPs with the highest connectivity to polyphenolic metabolites, providing insights into key genetic components influencing pigment biosynthesis. Partial correlation coefficients (*r*) and *P*-values were computed pairwise across all SNPs using the ppcor package (Kim 2015). SNP pairs with significant positive correlations (*r* = 0.5, *P* ≤ 0.05) were filtered and visualized as a correlation network in Cytoscape v3.10.1 (Shannon et al. 2003) to assess their potential roles in polyphenol-associated pathways.

### Database development

SorGMDA was developed using MySQL and PHP to enable users to search metabolome profiling data acquired from the SAP accessions studied. The database integrates mGWAS signals with *P*-values ≤ 10□³ mapped across all chromosomes.

## Supporting information

Revised_Supplementary_files.zip_1_to_12

## Data availability

All metabolic traits and corresponding GWAS results can be accessed through SorGMDA (https://www.depts.ttu.edu/igcast/SorbGMDA/index.php).

## Code availability

All analyses were performed using publicly available software and standard computational workflows as described in the Methods section. The full set of scripts used in this study is available in the Supplemental Material (Code S1–S3). Corresponding code is also openly accessible on GitHub as follows: https://github.com/Deepti098c/FilterVCF_Multilocus; https://github.com/Deepti098c/MetReg.plot; https://github.com/Deepti098c/sorghum_grain_color_ML.

## Author contributions

YJ conceptualized and designed the project. MY contributed to project discussions. NK, DN, RT and AK prepared samples for metabolomic analysis. SM, QS, and FC conducted the metabolome profiling. DN performed all data analyses. PK and DN designed the database. DN, YJ, and MY drafted the manuscript. All authors reviewed, edited, and approved the final version of the manuscript.

## Acknowledgments

This research was supported by the U.S. Department of Agriculture (USDA)’s Intramural Research Program (3096-21000-024-00D), the National Institute of Food and Agriculture’s Agriculture and Food Research Initiative (AFRI) under award number 2023-67013-39631, and the Non-Assistance Cooperative Agreement 58-3020-2-024 between Texas Tech University and USDA-ARS (3020-43440-002-00D). The authors are grateful to the publicly available USDA Germplasm Resources Information Network (GRIN) database for providing accessions of the SAP used in this study.

## Supplemental figures

**Supplemental Figure S1:**
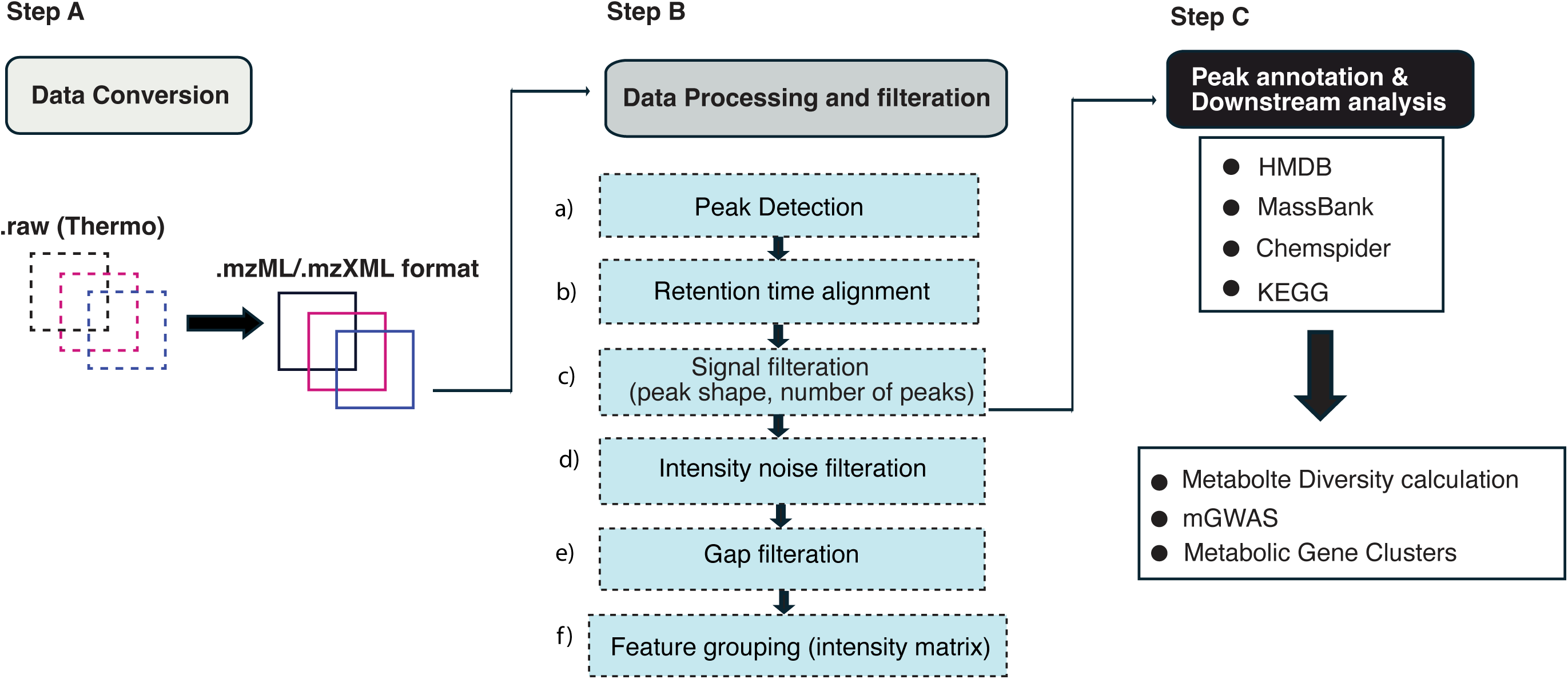
Workflow for mass spectrometry-based metabolomics analysis in the Sorghum Association Panel (SAP). The workflow for mass spectrometry-based metabolomics analysis covered data filtration, peak annotation, and downstream analyses. In Step A, raw data from Thermo Fisher Scientific mass spectrometers (.raw format) are converted to standardized formats (.mzML/.mzXML) for further processing. Step B details several stages of data processing: peak detection identifies metabolites by their mass-to-charge ratio (m/z) and intensity; retention time alignment ensures consistency across samples; signal filtration improves data quality by evaluating peak shape and count; intensity noise filtration removes low-intensity noise; gap filtration addresses missing data points; and feature grouping organizes the processed data into an intensity matrix. In Step C, processed peaks are annotated using established databases, such as HMDB, MassBank, ChemSpider, and KEGG. The workflow concludes with downstream analyses, including the calculation of metabolite diversity, mGWAS, and validation of results through further experimental or computational methods.

**Supplemental Figure S2.**
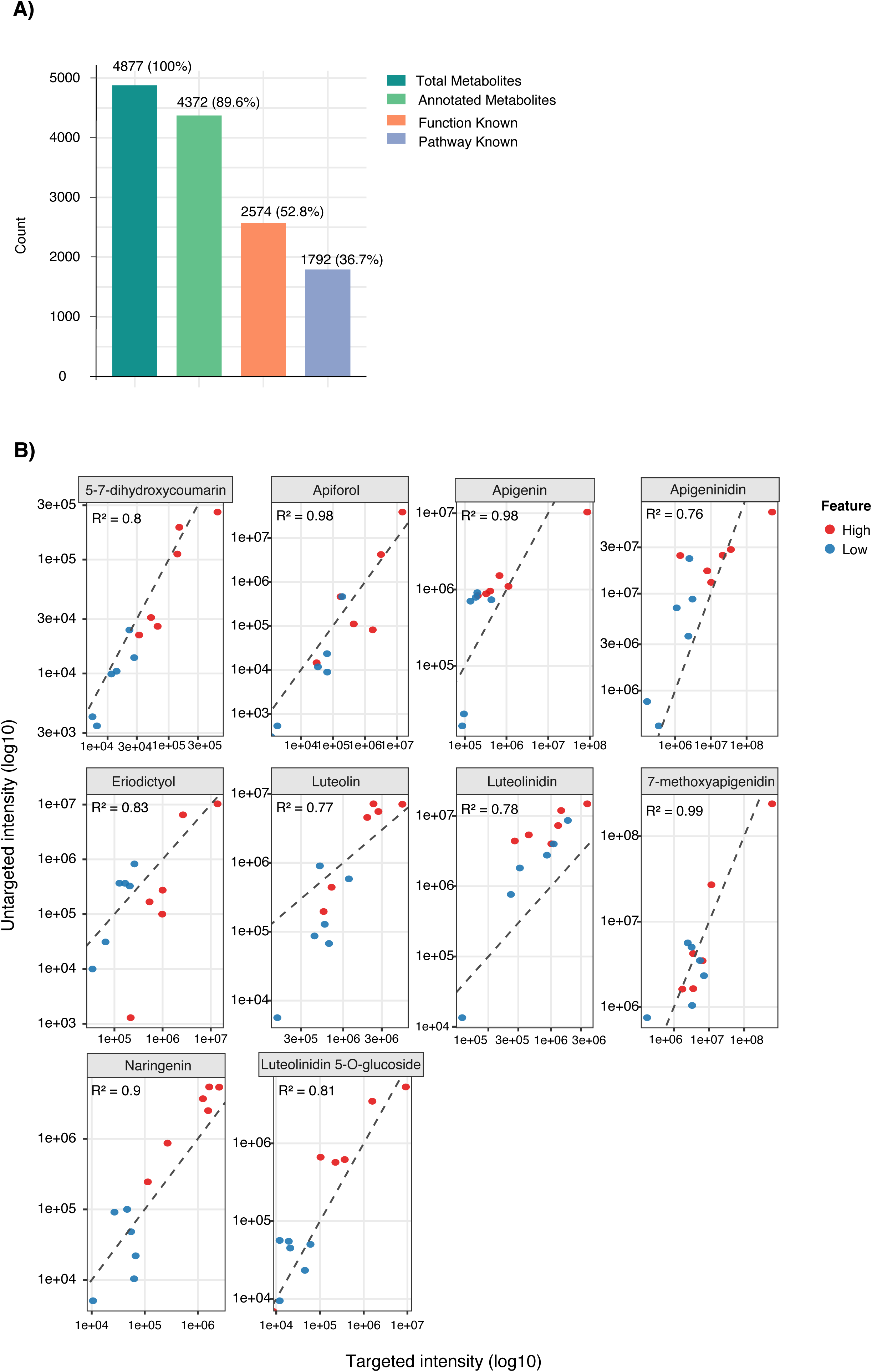
Annotation summary and validation of untargeted metabolite profiling in sorghum grain. (A) Summary of metabolite annotation statistics from untargeted LC–MS profiling of sorghum grain samples. Bars indicate the total number of detected metabolites, annotated metabolites, functionally characterized metabolites, and metabolites assigned to known metabolic pathways. Percentages relative to the total detected metabolites are shown above each bar. (B) Correlation analysis between untargeted and targeted metabolite quantification for representative polyphenol compounds, including flavonoids and phenolic acids. Scatter plots compare normalized untargeted LC–MS intensities with targeted metabolite measurements on a log10 scale. Dashed lines indicate linear regression fits, and corresponding Pearson correlation coefficients (R²) are shown for each compound. Red and blue points represent samples with relatively high and low metabolite abundance, respectively.

**Supplemental Figure S3.**
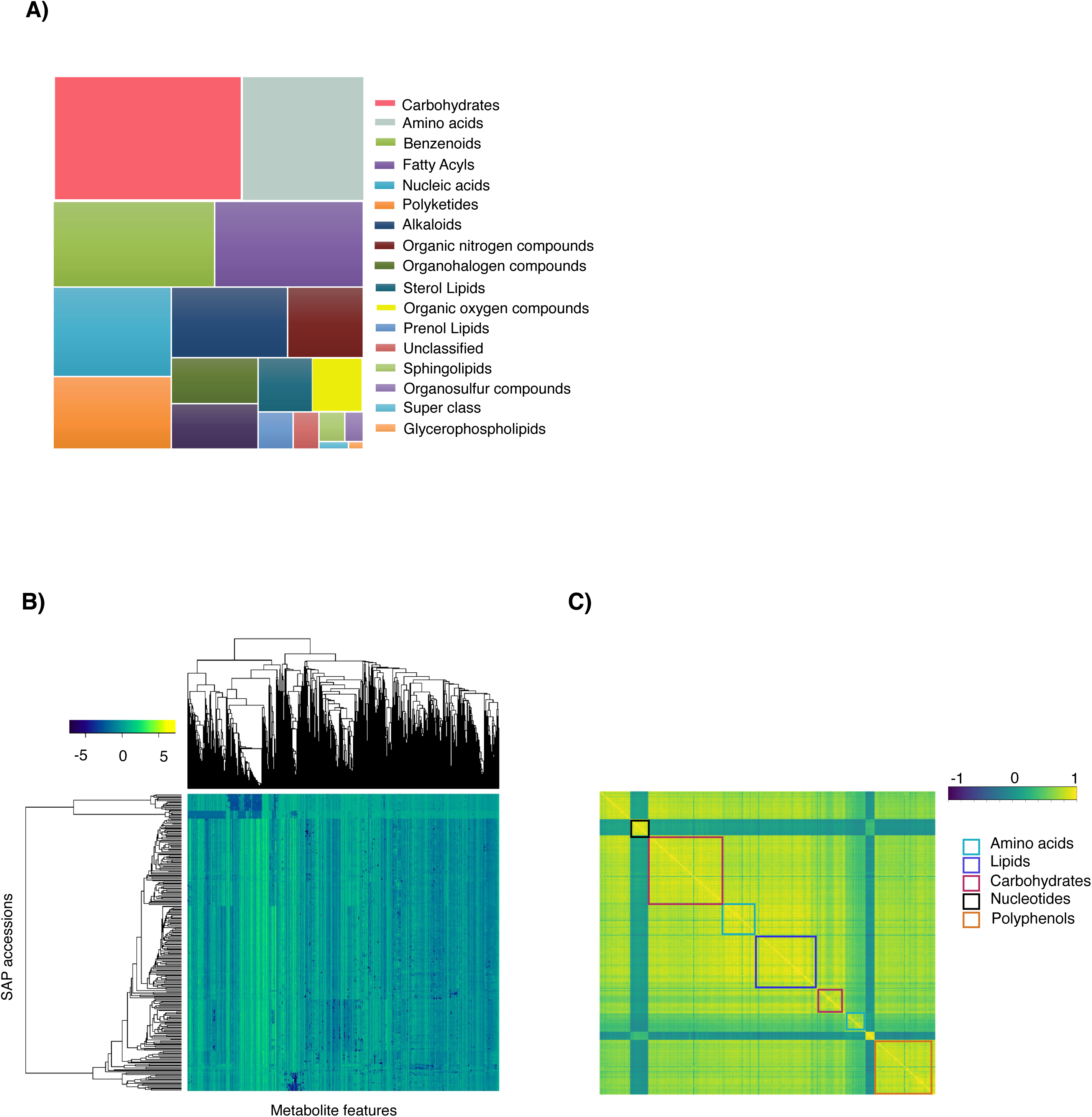
Functional categorization and correlation structure of identified metabolites in sorghum grain. (A) Treemap visualization showing the distribution of identified metabolites across major chemical superclasses. Each colored block represents a metabolite class, and block size is proportional to the number of metabolites assigned to that category. (B) Hierarchical clustering heatmap of metabolite abundance profiles across the 266 Sorghum Association Panel (SAP) accessions. Rows represent SAP accessions and columns represent metabolite features. Color intensity indicates relative metabolite abundance following normalization and scaling. Dendrograms illustrate similarity relationships among metabolites and accessions. (C) Correlation matrix of metabolite abundance profiles grouped by major functional categories, including amino acids, lipids, carbohydrates, nucleotides, and polyphenols. Colored boxes highlight clusters of metabolites exhibiting coordinated accumulation patterns, suggesting shared biochemical pathways or co-regulated metabolic processes.

**Supplemental Figure S4.**
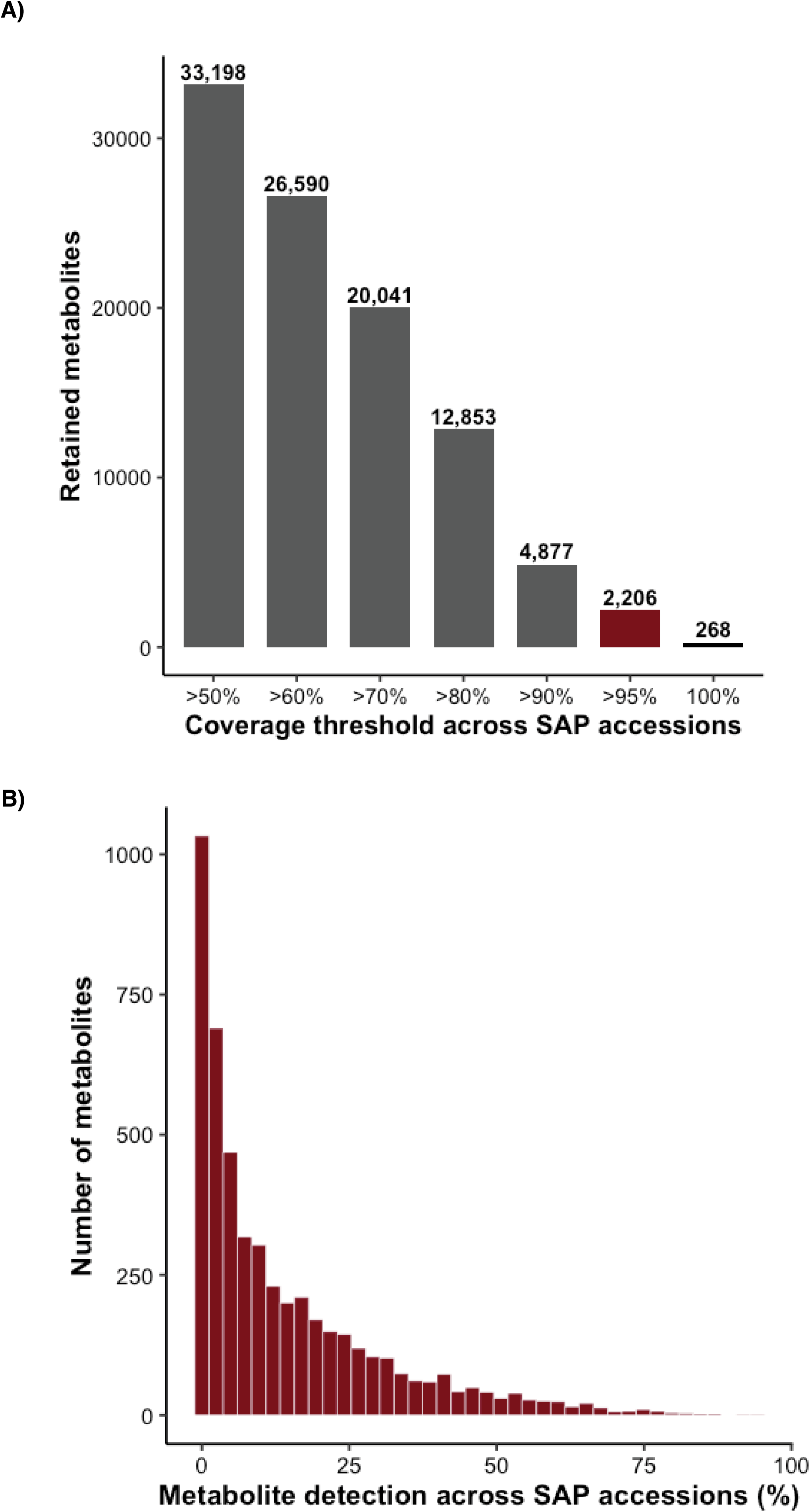
Detection frequency and coverage distribution of metabolites across the Sorghum Association Panel (SAP). (A) Number of metabolites retained at increasing population coverage thresholds across SAP accessions. A total of 2,206 metabolites detected in >95% of accessions were retained for downstream analyses. (B) Distribution of metabolite detection frequencies across SAP accessions, showing that most metabolites were detected in a subset of samples.

**Supplemental Figure S5.**
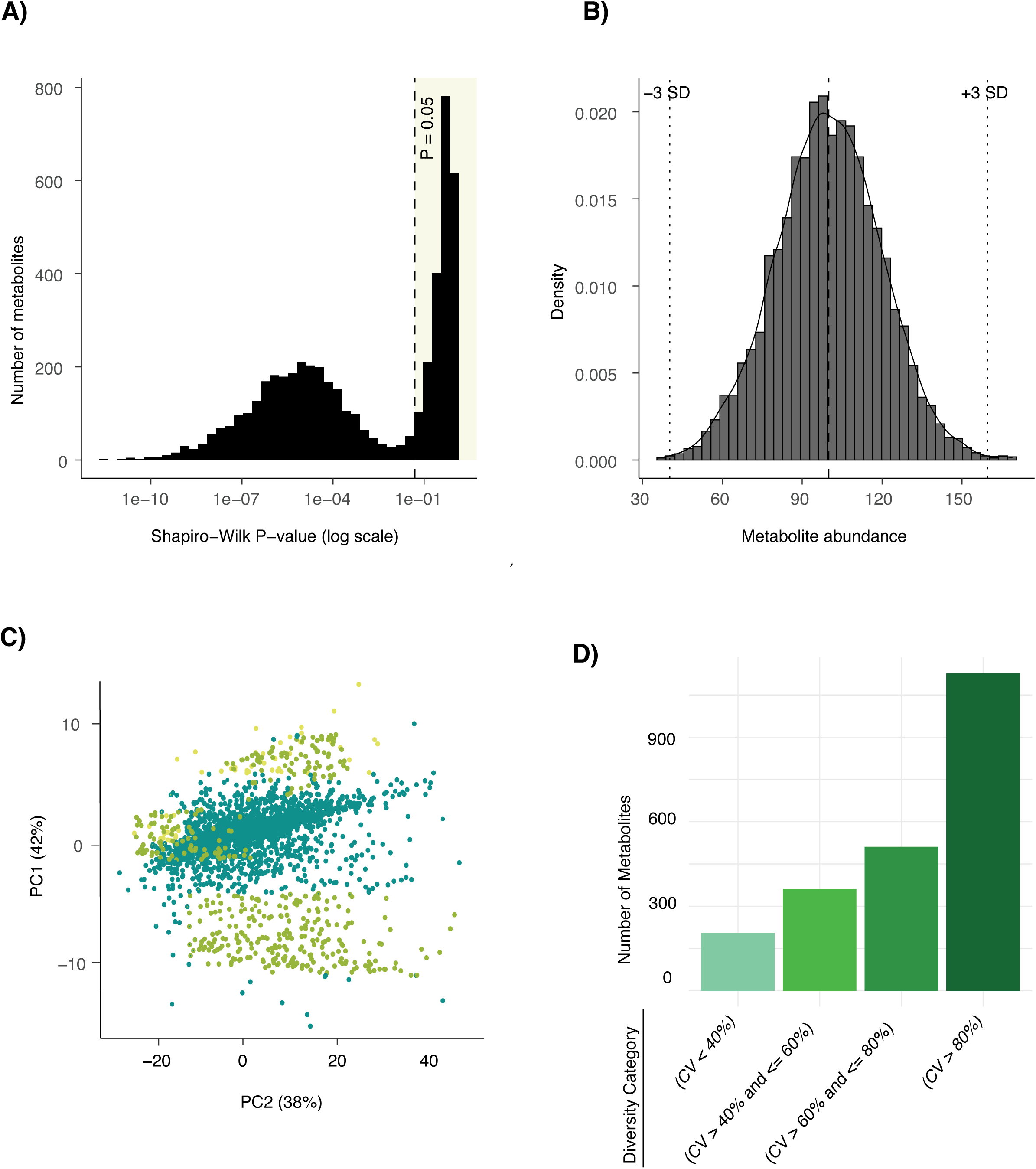
Quality control (QC) normalization and metabolite filtering prior to downstream analysis. (A) Distribution of Shapiro–Wilk test p-values for all detected metabolites after log transformation. The shaded region indicates metabolites passing the normality threshold (P > 0.05). (B) Representative distribution of normalized metabolite abundance values after QC normalization, with ±3 standard deviation (SD) thresholds used for outlier filtering. Dashed vertical lines indicate the lower and upper ±3 SD boundaries. (C) Principal component analysis (PCA) of retained metabolites, colored by cohort/AP group. (D) Distribution of metabolite coefficients of variation (CV) across samples, grouped into four variability categories: CV < 40%, CV = 40–60%, CV = 60–80%, and CV > 80%.

**Supplemental Figure S6.**
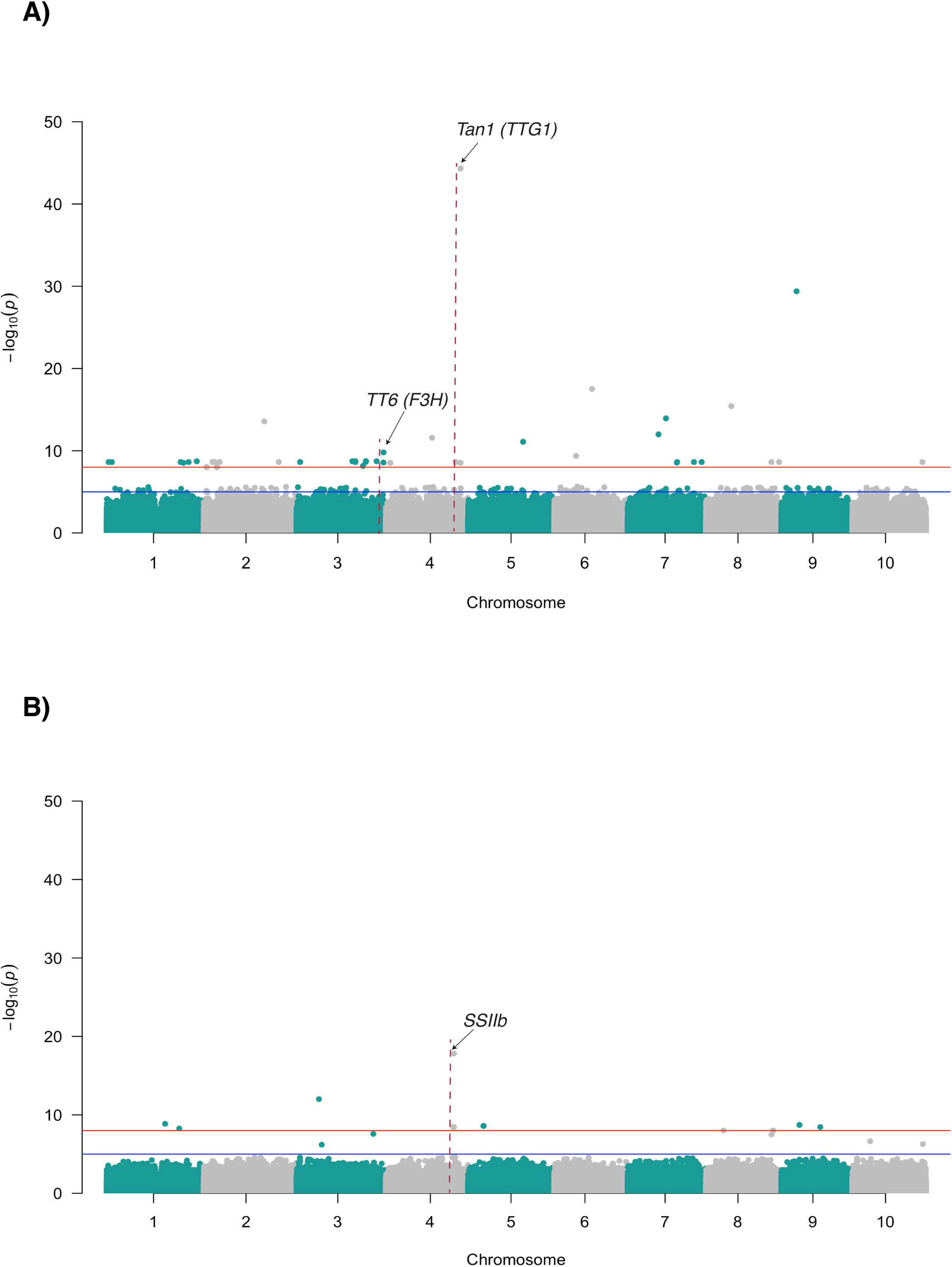
Genome-wide association signals for (−)-epigallocatechin and D-(−)-fructose. (A) Manhattan plot showing mGWAS results for (−)-epigallocatechin (C□□H□□O□). Significant association peaks were detected near the *Tan1 (TTG1)* and *TT6 (F3H)* loci. (B) Manhattan plot showing mGWAS results for D-(−)-fructose (C□H□□O□). A major association peak was detected near the SSIIb locus. In both panels, each point represents a SNP, with chromosomal position shown on the x-axis and statistical significance [−log10(P)] shown on the y-axis. Horizontal lines indicate significance thresholds.

**Supplemental Figure S7.**
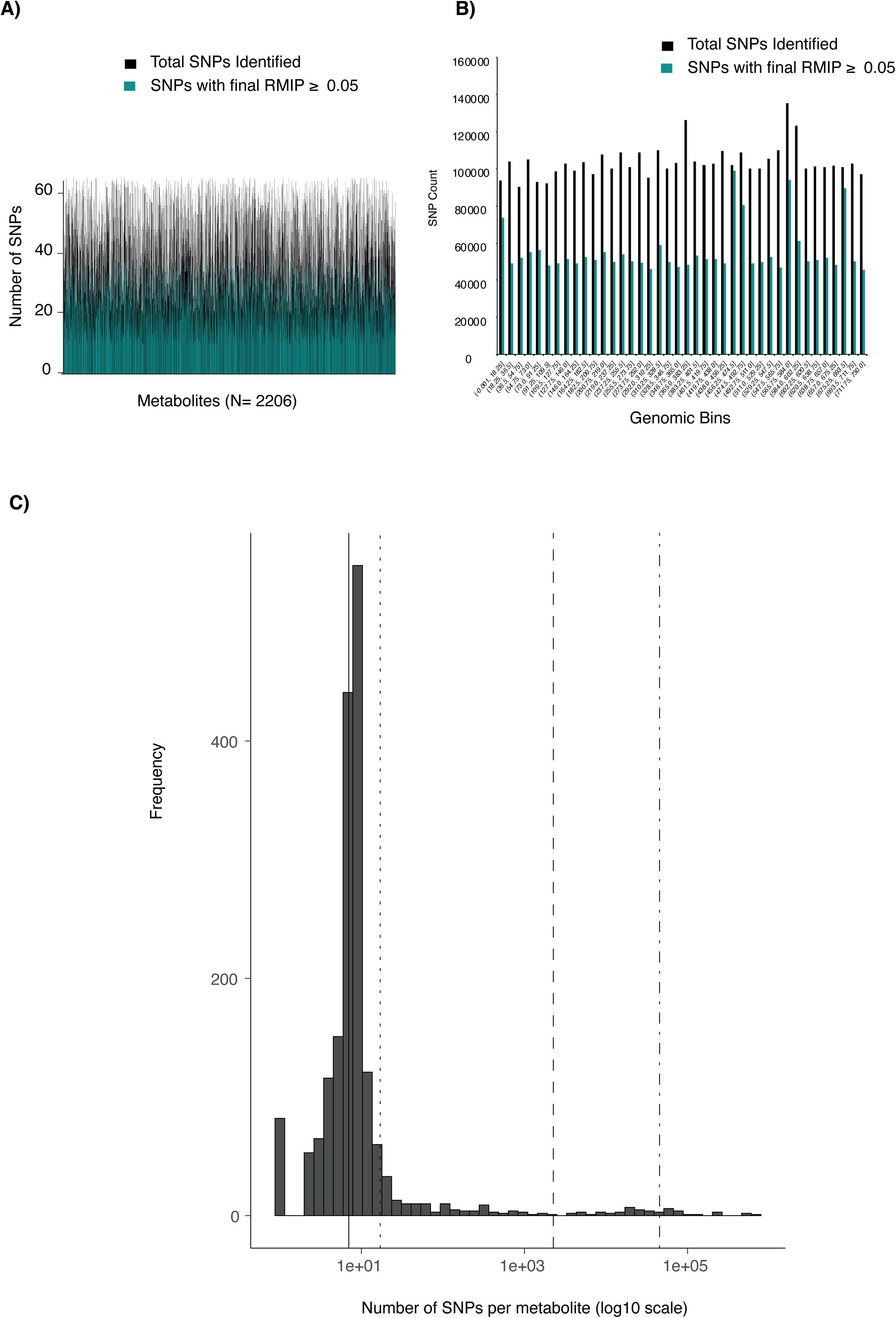
Distribution of metabolite values and SNP association prioritization. (A) Distribution of 2206 metabolite values demonstrating normality across samples (Shapiro-Wilk test P > 0.05; within ±3 standard deviations). The histogram displays the frequency of metabolite measurements, with an overlaid normal distribution curve, illustrating the retention of metabolites that exhibit normal distribution characteristics. (B) Comparison of the total number of SNPs identified per metabolite and the subset of SNPs with resample model inclusion probability (RMIP) ≥ 0.05. The RMIP threshold was used to prioritize SNPs consistently detected across multiple resampling iterations, enhancing the reliability and reproducibility of association signals; (C) Distribution of significant SNP–metabolite associations per metabolite. Histogram showing the number of significant SNPs associated with each of the 2,206 metabolites retained for mGWAS analysis. The x-axis is shown on a log10 scale. Vertical dashed lines indicate the median, 90th percentile, and top 1% of metabolites.

**Supplemental Figure S8.**
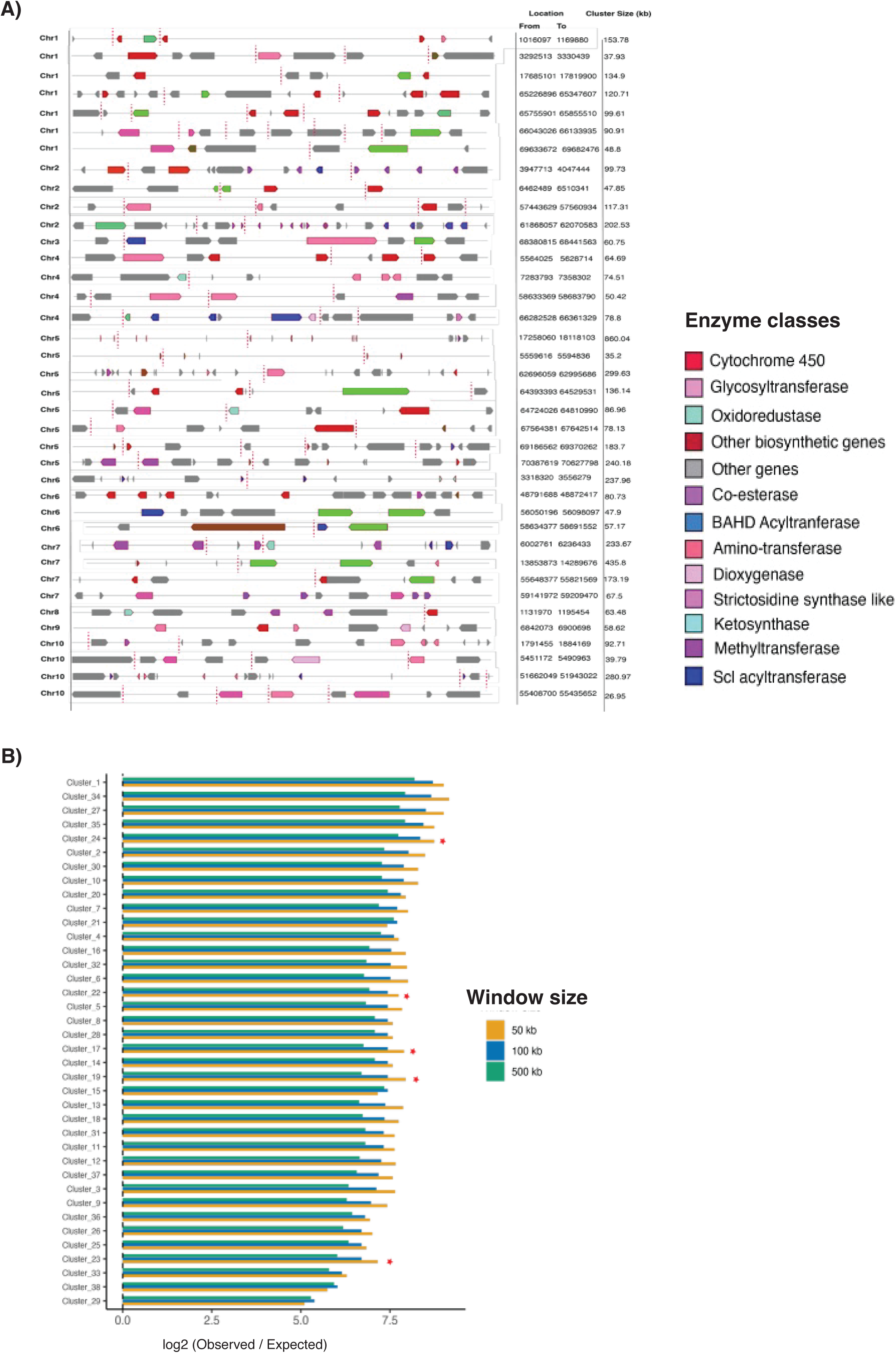
Genomic distribution of metabolic gene clusters and enrichment of metabolite-associated SNPs in sorghum. (A) Genome-wide distribution of 38 metabolic gene clusters (MGCs) identified in the Sorghum bicolor genome. Clusters are labeled sequentially from Cluster 1 to Cluster 38. Red dotted vertical lines indicate the positions of significant metabolite-associated SNPs identified through mGWAS. The right panel shows the genomic position and relative size of each metabolic gene cluster across chromosomes. (B) Cluster-wise enrichment of mGWAS SNP associations across metabolic gene clusters. Bar plots show enrichment of metabolite-associated SNPs within genomic windows surrounding each cluster (±50 kb, ±100 kb, and ±500 kb). Enrichment values are calculated as log□(observed/expected), where positive values indicate overrepresentation of significant SNPs relative to genome-wide expectations.

**Supplemental Figure S9.**
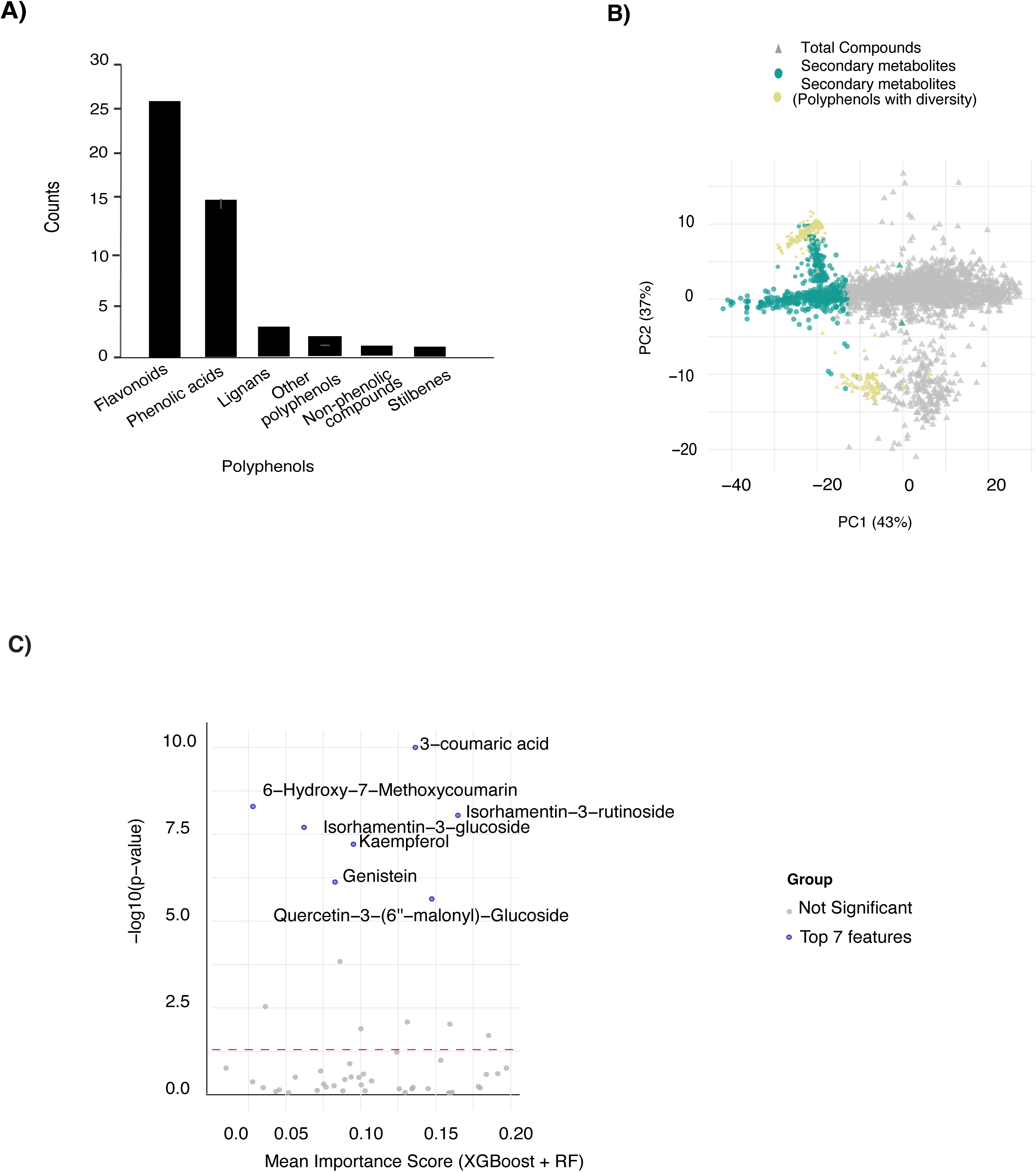
Grain polyphenol diversity and enzymatic pathway variation across SAP accessions. (A) 49 Polyphenols identified across four grain color groups. (B) PCA of metabolite profiles across 211 accessions. PC1 and PC2 explain 43% and 37% of the total variance, respectively. Accessions are color-coded by grain color groups: brown (77 lines), red (22 lines), yellow (10 lines), and white (102 lines). (C) Metabolites are shown as dots with grey representing all detected metabolites, green indicating secondary metabolites, and yellow highlighting 49 polyphenols exhibiting diversity across the SAP.

**Supplemental Figure S10.**
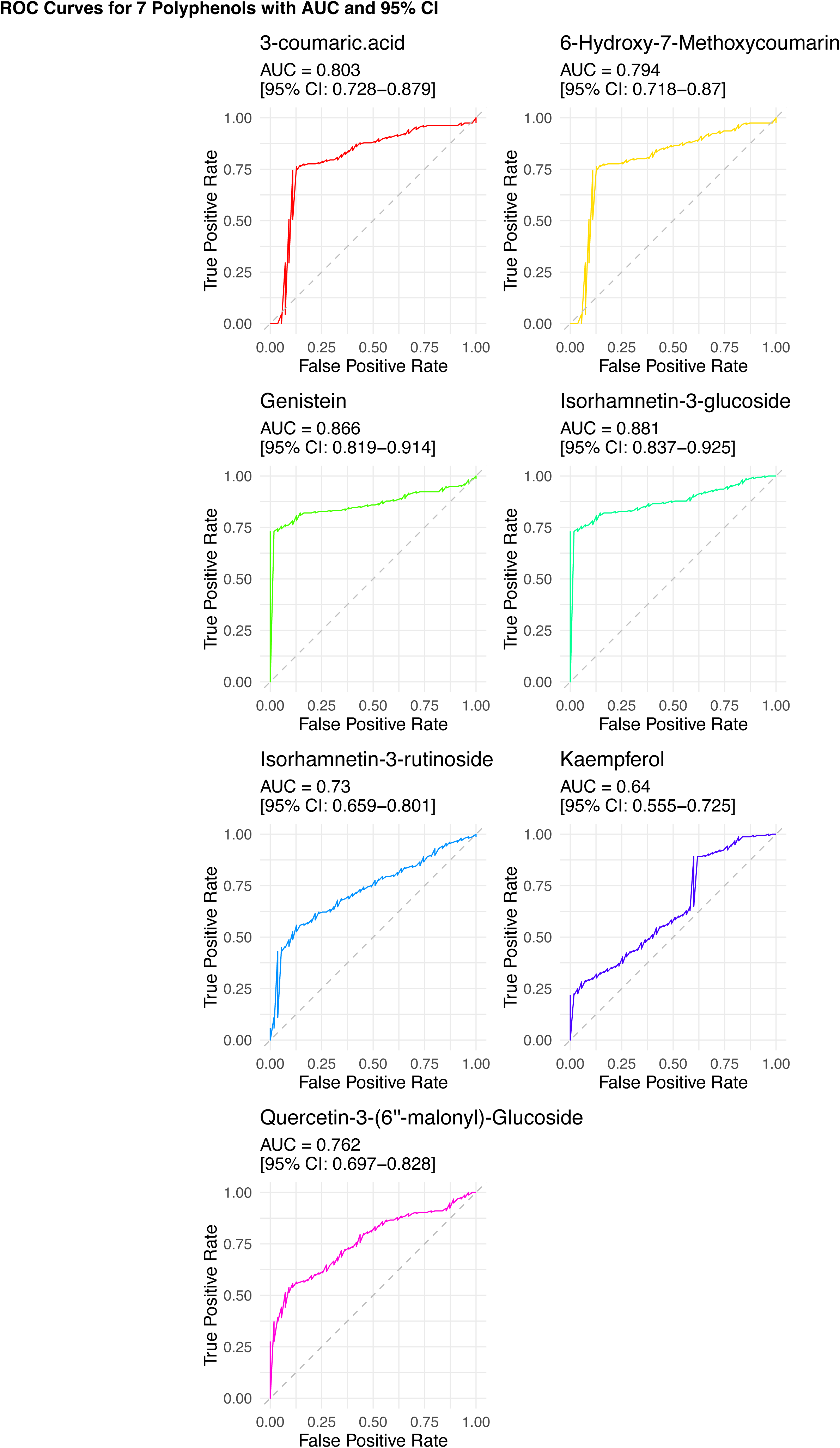
Receiver Operating Characteristic (ROC) curves for seven polyphenols showing their performance in distinguishing between 4 groups of sorghum accessions with distinct grain color. Each panel displays the ROC curve for an individual polyphenol, with the shaded area representing the 95% confidence interval around the curve. The diagonal dashed line indicates the performance of a random classifier (AUC = 0.5). These curves illustrate the discriminatory power of individual compounds based on their true-positive and false-positive rates.

**Supplemental Figure S11.**
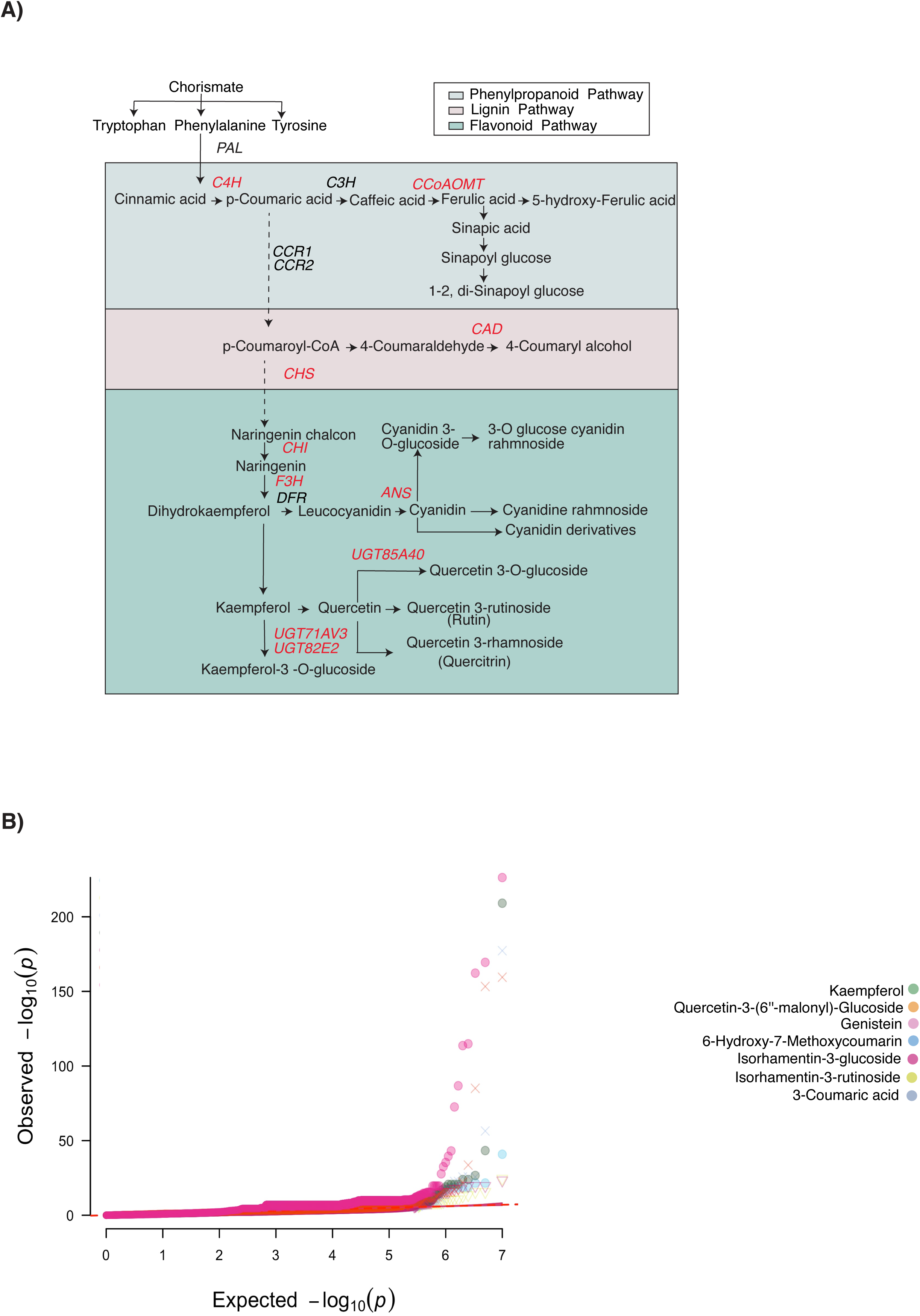
Biosynthetic pathway mapping and GWAS signal validation for key polyphenol-associated loci in sorghum. (A) Schematic representation of the phenylpropanoid, flavonoid, and lignin biosynthetic pathways. Enzymes detected near regions of genetic variation (±50 kb) are indicated, linking metabolite diversity with potential regulatory loci. (B) Quantile-quantile (QQ) plot of GWAS results for seven selected flavonoids. The plot displays the observed versus expected −log□□(p) values for SNP associations with metabolite abundances. Each color represents a different metabolite. The deviation from the diagonal red line indicates SNPs with stronger-than-expected associations, highlighting potential genetic loci controlling metabolite variation.

**Supplemental Figure S12.**
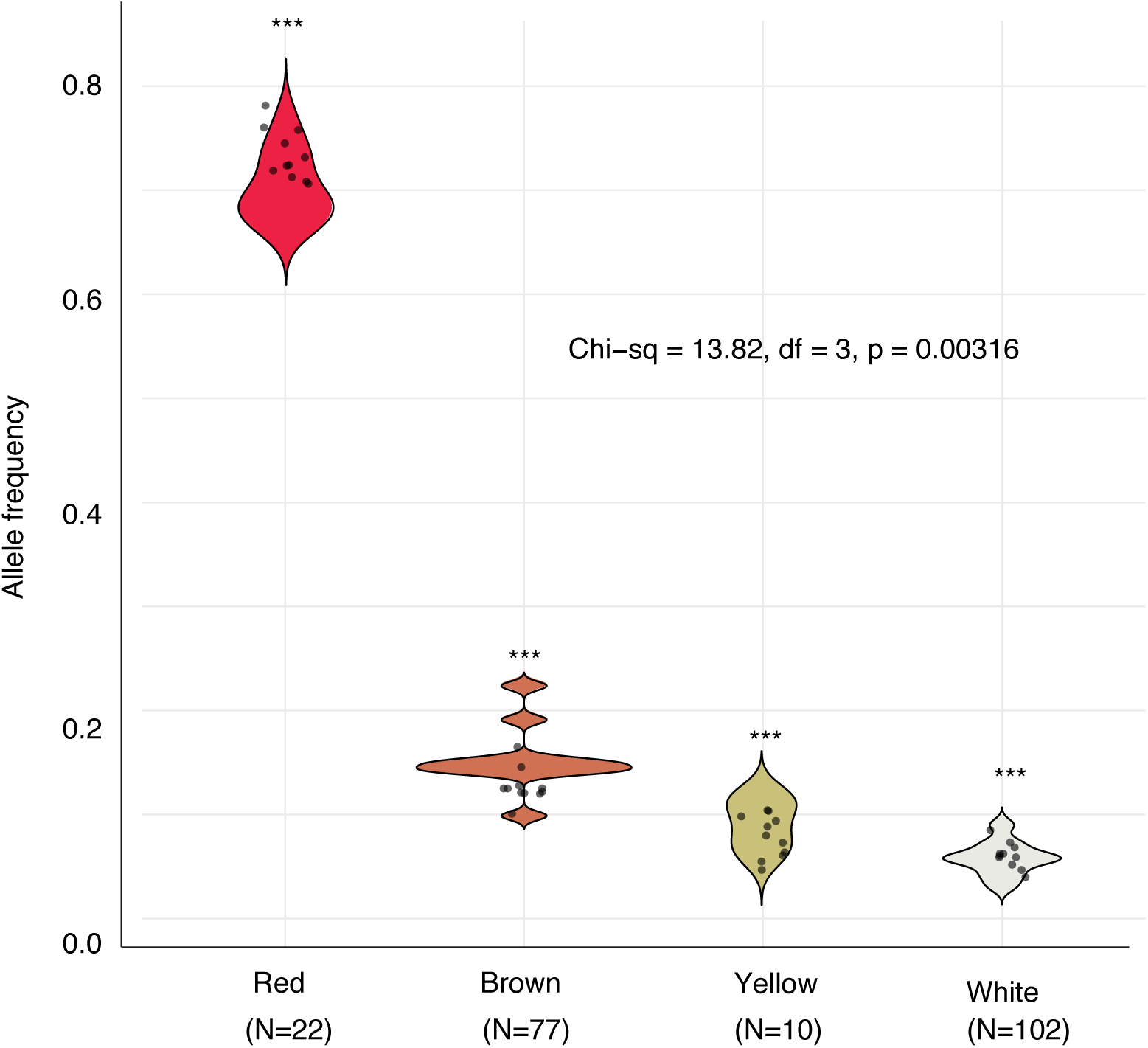
Allele frequency distribution of the high-polyphenol allele group across four grain color classes. Violin plots show the distribution of allele frequencies across red, brown, yellow, and white grain color classes. High-polyphenol alleles were predominantly enriched in darker grains (red and brown) and occurred at substantially lower frequencies in lighter grains (yellow and white). A chi-square test confirmed a significant association between allele frequency and grain color class (χ² = 13.82, df = 3, *P* = 0.00316).

**Supplemental Figure S13.**
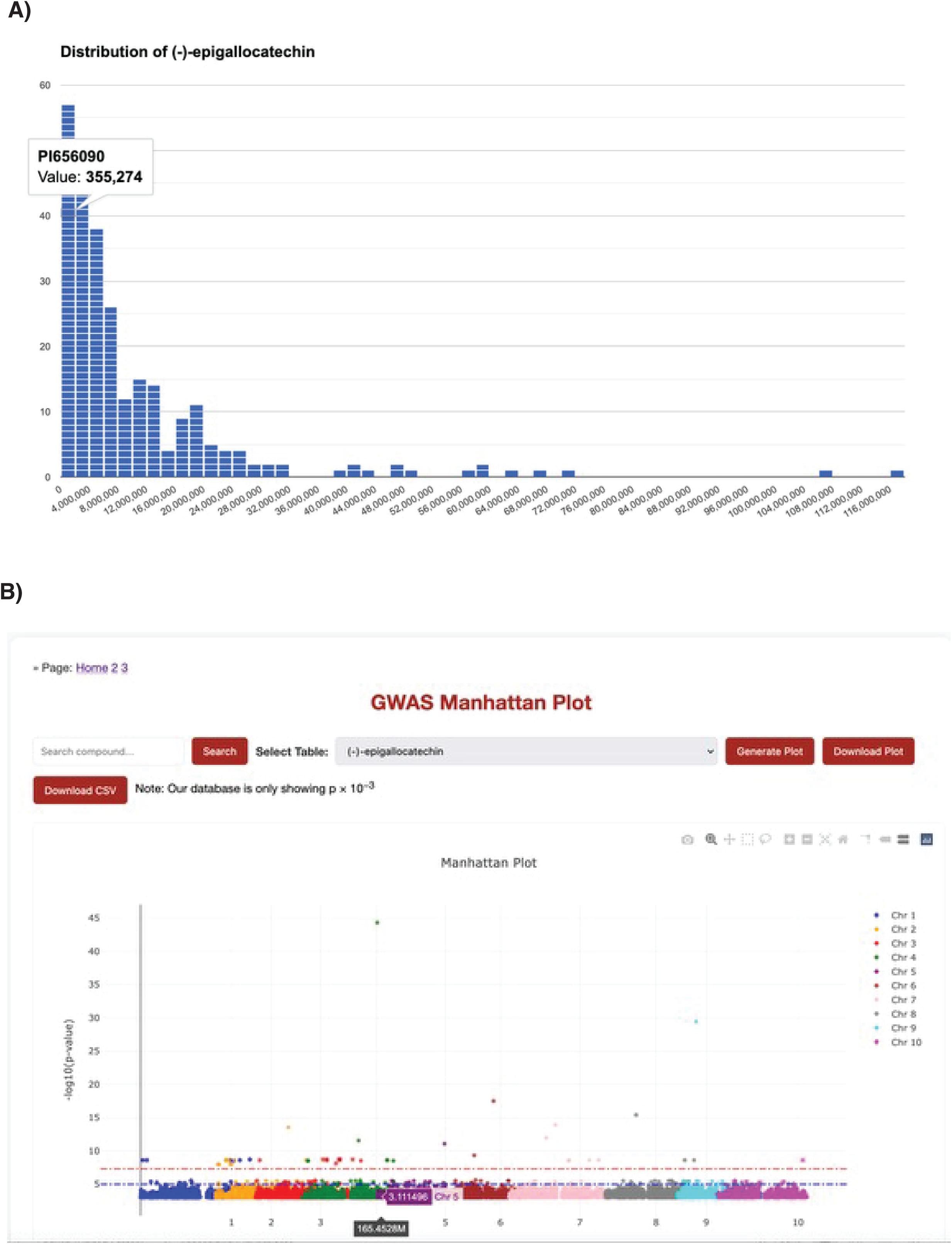
Natural variation and genome-wide association mapping of (−)-epigallocatechin in sorghum seeds. (A) Distribution of (-)-epigallocatechin across GRIN-based SAP line numbers, with corresponding abundance values measured by UPLC-MS, obtained using SorGMDA. (B) Manhattan plot of genome-wide association study (GWAS) results for (–)-epigallocatechin accumulation in the SAP population. The plot displays -log□□(p-value) for each SNP across all sorghum chromosomes (Chr 1–10), with alternating colors used to distinguish between chromosomes. Only associations with p-values ≤ 10□³ are shown for visualization purposes; therefore, the number of plotted points may differ from the full GWAS dataset presented in Figure S4. Each dot represents a single nucleotide polymorphism (SNP), with the vertical axis indicating the statistical significance of association and the horizontal axis representing genomic position. A strong association signal is observed on chromosome 3, with a peak at position 165.4528 Mb showing a –log□□(p-value) of 45. The red and blue dashed lines represent suggestive and genome-wide significance thresholds, respectively.

## Supplemental Table

**Table S1.** Consensus polyphenolic features identified by XGBoost and Random Forest models for discriminating grain color groups. Note: Importance scores were normalized to represent the relative contribution of each compound in distinguishing grain pigment groups. Adjusted p-values were calculated using one-way ANOVA followed by Benjamini–Hochberg false discovery rate (FDR) correction for multiple testing. The −log10(p-value) metric was used for visualization, where higher values indicate greater statistical significance.

| Polyphenolic Feature | XGBoost Score | RF Score | Mean Importance | p-value | $-\log_{10}(\text{p-value})$ | Consensus Feature |
| --- | --- | --- | --- | --- | --- | --- |
| 3-coumaric acid | 0.084 | 0.072 | 0.078 | 0.002 | 2.7 | Yes |
| 6-Hydroxy-7-Methoxycoumarin | 0.066 | 0.054 | 0.06 | 0.015 | 1.82 | Yes |
| Genistein | 0.091 | 0.077 | 0.084 | 0.001 | 3 | Yes |
| Isorhamnetin-3-glucoside | 0.074 | 0.062 | 0.068 | 0.008 | 2.1 | Yes |
| Isorhamnetin-3-rutinoside | 0.079 | 0.069 | 0.074 | 0.009 | 2.05 | Yes |
| Kaempferol | 0.096 | 0.083 | 0.09 | 0.0005 | 3.3 | Yes |
| Quercetin-3-(6"-malonyl)-Glucoside | 0.088 | 0.071 | 0.08 | 0.003 | 2.52 | Yes |

**Note:** Importance scores were normalized to represent the relative contribution of each compound in distinguishing grain pigment groups. Adjusted p-values were calculated using one-way ANOVA followed by Benjamini–Hochberg false discovery rate (FDR) correction for multiple testing. The $-\log_{10}(\text{p-value})$ metric was used for visualization, where higher values indicate greater statistical significance.

## Supplemental files

**Supplemental Table S1.** Table listing the metabolite features detected via LC-MS, including Feature ID, m/z (mass-to-charge ratio), retention time (rtime), molecular formula, and compound name. Peak intensities are shown for each sorghum accession (PIXXXXXX), representing relative abundance values derived from the metabolomics dataset.

**Supplemental Table S2.** Comparative validation of targeted and untargeted metabolite measurements for 10 flavonoid and phenylpropanoid compounds using authentic standards.

**Supplemental Table S3.** List of metabolite features annotated with mass-to-charge ratio (m/z), retention time (rtime), molecular formula, and compound name. Each metabolite is characterized by its coefficient of variation (CV), diversity classification based on CV thresholds, and repeatability. Diversity categories are defined as follows: Low (CV < 40%), Medium (CV 40–80%), and High (CV > 80%).

**Supplemental Table S4:** List of candidate genes located within the ±50 kb genomic window of lead SNPs associated with the accumulation of (-)-epigallocatechin (C□□H□□O□), D-(-)-fructose (C□H□□O□), and lysine (C_6_H_14_N_2_O_2_) metabolites, identified through GWAS and supported by functional annotation. The table includes metabolite names, gene identifiers, chromosomal locations, SNP positions, gene boundaries (start and end coordinates), p-values, and corresponding gene annotations. Genes annotated as "putative" or "uncharacterized" reflect limited or predicted functional information as currently available in the Phytozome database (version 14).

**Supplemental Table S5.** Summary of SNP discovery and binning across ∼18 Mb genomic intervals for ten selected metabolites. The table includes compound names, total SNPs identified, SNPs with a final Resample Model Inclusion Probability (RMIP) > 0.05, genomic bins (each approximately 18 Mb in size), total SNP counts per bin, and the count of SNPs within each bin that meet the RMIP threshold. RMIP values reflect the robustness of SNP-trait associations across resampling iterations in genome-wide association models.

**Supplemental Table S6.** Comparison of the genomic distribution of SNPs identified through mGWAS versus the total input SNPs used in the analysis. Genomic regions include 3’ and 5’ UTRs, upstream and downstream regulatory regions, intergenic areas, introns, missense, and synonymous coding sites. Counts and percentages reflect the proportion of SNPs within each genomic category out of the total SNPs detected in mGWAS (left) and the full input SNP set (right). This analysis highlights the genomic context of SNP-trait associations and their relative enrichment in functional or regulatory regions.

**Supplemental Table S7.** List of 83 genes located within or near Metabolic Gene Clusters (MGCs) in the sorghum genome that harbor genomic variants. The table includes gene names, chromosomal coordinates, variant site locations, distances between variants and gene boundaries (in kb), and corresponding GWAS p-values. Each gene is annotated with available functional information including Gene Ontology (GO), PFAM, InterPro, KEGG, KOG, Panther, and ENZYME classifications.

**Supplemental Table S8.** Normalized expression profiles of lignin-associated genes during seed development in *Sorghum bicolor* across 45 RNA-seq samples. Samples are grouped into early whole-seed (se1–se9), embryo (em10–em25), and endosperm (en6–en25) developmental stages. Expression values represent the relative transcript abundance of each gene across developmental time points and tissues. Functional annotations were assigned based on KEGG and other curated pathway databases. A value of zero indicates that no detectable expression was observed in the corresponding sample.

**Supplemental Table S9.** Classification of SAP accessions based on four grain color groups (white, red, brown, and yellow), as recorded in the Germplasm Resources Information Network (GRIN) database.

**Supplemental Table S10.** Secondary metabolites associated with sorghum seed color variation. Summary of 49 secondary metabolites identified in sorghum seeds that showed significant associations with seed color variation among accessions from the Sorghum Association Panel (SAP). Sorghum accessions were classified into four seed color groups: white, red, brown, and yellow. For each metabolite, the table includes normalized importance scores, adjusted P-values, and corresponding −log10(P-value) values to facilitate interpretation of statistical significance. Quantitative values represent normalized metabolite abundances across samples within each seed color group.

**Supplemental Table S11.** SAP accessions, grain color category (white, red, brown, and yellow), binary true labels, and normalized abundances for seven key metabolites associated with grain color variation. Metabolite values represent relative abundances normalized across all samples. The true labels column indicates binary classification (e.g., presence or absence of a trait) used in predictive modeling or phenotype association analyses.

**Supplemental Table S12.** Metabolites and genetic mapping results identifying candidate genes associated with variation among four grain color groups. For each metabolite, the table lists the statistical model(s) used for association analysis (FarmCPU, MLM), chromosome number, standardized effect size, lead SNP position (in base pairs), p-value of association, mapped gene ID, gene start and end positions, gene annotation, and reference genome version (B.Tx623 v3.1).

## References

Ali AEE, Sharp RE, Greeley L, Peck SC, Tabb DL, Ludidi N. 2025. Proteomic dataset of sorghum leaf and root responses to single and combined drought and heat stress. Scientific Data 12: 97.

Almodares A, Hadi M. 2009. Production of bioethanol from sweet sorghum: A review. African journal of agricultural research 4: 772–780.

Arbex PM, de Castro Moreira ME, Toledo RCL, de Morais Cardoso L, Pinheiro-Sant’ana HM, dos Anjos Benjamin L, Licursi L, Carvalho CWP, Queiroz VAV, Martino HSD. 2018. Extruded sorghum flour (Sorghum bicolor L.) modulate adiposity and inflammation in high fat diet-induced obese rats. Journal of functional foods 42: 346-355.

Aron AT, Gentry EC, McPhail KL, Nothias L-F, Nothias-Esposito M, Bouslimani A, Petras D, Gauglitz JM, Sikora N, Vargas F. 2020. Reproducible molecular networking of untargeted mass spectrometry data using GNPS. Nature protocols 15: 1954–1991.

Awika JM, Rooney LW. 2004. Sorghum phytochemicals and their potential impact on human health. Phytochemistry 65: 1199–1221.

Awika JM, Yang L, Browning JD, Faraj A. 2009. Comparative antioxidant, antiproliferative and phase II enzyme inducing potential of sorghum (Sorghum bicolor) varieties. LWT-Food Science and Technology 42: 1041–1046.

Baniasadi H, Vlahakis C, Hazebroek J, Zhong C, Asiago V. 2014. Effect of environment and genotype on commercial maize hybrids using LC/MS-based metabolomics. Journal of Agricultural and Food Chemistry 62: 1412–1422.

Bi B, Wang K, Zhang H, Wang Y, Fei H, Pan R, Han F. 2021. Plants use rhizosphere metabolites to regulate soil microbial diversity. Land Degradation & Development 32: 5267–5280.

Bian Y, Holland J. 2017. Enhancing genomic prediction with genome-wide association studies in multiparental maize populations. Heredity 118: 585–593.

Boatwright JL, Sapkota S, Jin H, Schnable JC, Brenton Z, Boyles R, Kresovich S. 2022. Sorghum Association Panel wholecgenome sequencing establishes cornerstone resource for dissecting genomic diversity. The Plant Journal 111: 888–904.

Bordiga M, Gomez-Alonso S, Locatelli M, Travaglia F, Coïsson JD, Hermosin-Gutierrez I, Arlorio M. 2014. Phenolics characterization and antioxidant activity of six different pigmented Oryza sativa L. cultivars grown in Piedmont (Italy). Food Research International 65: 282–290.

Boyles RE, Pfeiffer BK, Cooper EA, Rauh BL, Zielinski KJ, Myers MT, Brenton Z, Rooney WL, Kresovich S. 2017. Genetic dissection of sorghum grain quality traits using diverse and segregating populations. Theoretical and applied genetics 130: 697–716.

Bröhan M, Jerkovic V, Wilmotte R, Collin S. 2011. Catechins and derived procyanidins in red and white sorghum: Their contributions to antioxidant activity. Journal of the Institute of Brewing 117: 600–607.

Bunsupa S, Yamazaki M, Saito K. 2017. Lysine-derived alkaloids: overview and update on biosynthesis and medicinal applications with emphasis on quinolizidine alkaloids. Mini Reviews in Medicinal Chemistry 17: 1002–1012.

Burdette A, Garner PL, Mayer EP, Hargrove JL, Hartle DK, Greenspan P. 2010. Anti-inflammatory activity of select sorghum (Sorghum bicolor) brans. Journal of medicinal food 13: 879–887.

Burke HB. 2023. Use the receiver operating characteristic to assess model accuracy. JAMA cardiology 8: 998–998.

Bursać M, Krstonošić MA, Miladinović J, Malenčić Đ, Gvozdenović L, Cvejić JH. 2017. Isoflavone composition, total phenolic content and antioxidant capacity of soybeans with colored seed coat. Natural Product Communications 12: 1934578X1701200417.

Buzas MA, Hayek L-AC. 2005. On richness and evenness within and between communities. Paleobiology 31: 199–220.

Byrne PF, Volk GM, Gardner C, Gore MA, Simon PW, Smith S. 2018. Sustaining the future of plant breeding: The critical role of the USDAcARS National Plant Germplasm System. Crop Science 58: 451–468.

Campbell BC, Gilding EK, Mace ES, Tai S, Tao Y, Prentis PJ, Thomelin P, Jordan DR, Godwin ID. 2016. Domestication and the storage starch biosynthesis pathway: signatures of selection from a whole sorghum genome sequencing strategy. Plant biotechnology journal 14: 2240–2253.

Carreno-Quintero N, Bouwmeester HJ, Keurentjes JJ. 2013. Genetic analysis of metabolome–phenotype interactions: from model to crop species. Trends in Genetics 29: 41–50.

Chaleckis R, Meister I, Zhang P, Wheelock CE. 2019. Challenges, progress and promises of metabolite annotation for LC–MS-based metabolomics. Current opinion in biotechnology 55: 44–50.

Chambers MC, Maclean B, Burke R, Amodei D, Ruderman DL, Neumann S, Gatto L, Fischer B, Pratt B, Egertson J. 2012. A cross-platform toolkit for mass spectrometry and proteomics. Nature biotechnology 30: 918–920.

Chen B-R, Wang C-Y, Ping W, Zhu Z-X, Ning X, Shi G-S, Miao Y, Nai W, Li J-H, Hou J-M. 2019. Genome-wide association study for starch content and constitution in sorghum (Sorghum bicolor (L.) Moench). Journal of Integrative Agriculture 18: 2446–2456.

Chen J, Hu X, Shi T, Yin H, Sun D, Hao Y, Xia X, Luo J, Fernie AR, He Z. 2020. Metabolitecbased genomecwide association study enables dissection of the flavonoid decoration pathway of wheat kernels. Plant Biotechnology Journal 18: 1722–1735.

Chen J, Xue M, Liu H, Fernie AR, Chen W. 2021a. Exploring the genic resources underlying metabolites through mGWAS and mQTL in wheat: From large-scale gene identification and pathway elucidation to crop improvement. Plant Communications 2.

Chen L, Zhang T, Li T. 2017. Gradient boosting model for unbalanced quantitative mass spectra quality assessment. In 2017 International conference on security, pattern analysis, and cybernetics (SPAC), pp. 394-399. IEEE.

Chen W, Wang W, Peng M, Gong L, Gao Y, Wan J, Wang S, Shi L, Zhou B, Li Z. 2016. Comparative and parallel genome-wide association studies for metabolic and agronomic traits in cereals. Nature communications 7: 12767.

Chen X, Shen J, Xu J, Herald T, Smolensky D, Perumal R, Wang W. 2021b. Sorghum phenolic compounds are associated with cell growth inhibition through cell cycle arrest and apoptosis in human hepatocarcinoma and colorectal adenocarcinoma cells. Foods 10: 993.

Cheng K, Dong R, Pan F, Su W, Xi L, Zhang M, Geng J, Gao R, Jin W, Abd El-Aty A. 2025. Characterization and feature selection of volatile metabolites in Yangxian pigmented rice varieties through GC-MS and machine learning algorithms. Frontiers in nutrition 12: 1598875.

Cingoz GS, Gurel E. 2016. Effects of salicylic acid on thermotolerance and cardenolide accumulation under high temperature stress in Digitalis trojana Ivanina. Plant Physiology and Biochemistry 105: 145–149.

Coklar H, Akbulut M. 2021. Changes in phenolic acids, flavonoids, anthocyanins, and antioxidant activities of Mahonia aquifolium berries during fruit development and elucidation of the phenolic biosynthetic pathway. Horticulture, Environment, and Biotechnology 62: 785–794.

Collins A, Santhakumar AB, Francis N, Blanchard C, Chinkwo K. 2024. Impact of sorghum (Sorghum bicolor L. Moench) phenolic compounds on cancer development pathways. Food Bioscience: 104177.

Council NR, Resources CoMGG, Imperatives A. 1990. The US national plant germplasm system.

DeJong TM. 1975. A comparison of three diversity indices based on their components of richness and evenness. Oikos: 222–227.

Desmet S, Saeys Y, Verstaen K, Dauwe R, Kim H, Niculaes C, Fukushima A, Goeminne G, Vanholme R, Ralph J. 2021. Maize specialized metabolome networks reveal organ-preferential mixed glycosides. Computational and Structural Biotechnology Journal 19: 1127–1144.

Domingo-Almenara X, Montenegro-Burke JR, Benton HP, Siuzdak G. 2017. Annotation: a computational solution for streamlining metabolomics analysis. Analytical chemistry 90: 480.

Dowd PF, Funnell-Harris DL, Sattler SE. 2016. Field damage of sorghum (Sorghum bicolor) with reduced lignin levels by naturally occurring insect pests and pathogens. Journal of Pest Science 89: 885–895.

Dykes L, Rooney WL, Rooney LW. 2013. Evaluation of phenolics and antioxidant activity of black sorghum hybrids. Journal of Cereal Science 58: 278–283.

Dykes L, Seitz LM, Rooney WL, Rooney LW. 2009. Flavonoid composition of red sorghum genotypes. Food Chemistry 116: 313–317.

Eiler J, Cesar J, Chimiak L, Dallas B, Grice K, Griep-Raming J, Juchelka D, Kitchen N, Lloyd M, Makarov A. 2017. Analysis of molecular isotopic structures at high precision and accuracy by Orbitrap mass spectrometry. International Journal of Mass Spectrometry 422: 126–142.

Erban A, Fehrle I, Martinez-Seidel F, Brigante F, Más AL, Baroni V, Wunderlin D, Kopka J. 2019. Discovery of food identity markers by metabolomics and machine learning technology. Scientific Reports 9: 9697.

Falcone Ferreyra ML, Emiliani J, Rodriguez EJ, Campos-Bermudez VA, Grotewold E, Casati P. 2015. The identification of maize and Arabidopsis type I flavone synthases links flavones with hormones and biotic interactions. Plant Physiology 169: 1090–1107.

Farrar JL, Hartle DK, Hargrove JL, Greenspan P. 2008. A novel nutraceutical property of select sorghum (Sorghum bicolor) brans: inhibition of protein glycation. Phytotherapy Research 22: 1052–1056.

Fernie AR, Schauer N. 2009. Metabolomics-assisted breeding: a viable option for crop improvement? Trends in genetics 25: 39–48.

Fiehn O, Wohlgemuth G, Scholz M, Kind T, Lee DY, Lu Y, Moon S, Nikolau B. 2008. Quality control for plant metabolomics: reporting MSIccompliant studies. The Plant Journal 53: 691–704.

Fontanari T, Fróes TC, Recamonde-Mendoza M. 2022. Cross-validation strategies for balanced and imbalanced datasets. In Brazilian Conference on Intelligent Systems, pp. 626-640. Springer.

Frei M. 2013. Lignin: characterization of a multifaceted crop component. The Scientific World Journal 2013: 436517.

Fukushima A, Kusano M, Redestig H, Arita M, Saito K. 2011. Metabolomic correlation-network modules in Arabidopsis based on a graph-clustering approach. BMC systems biology 5: 1.

Galili G, Amir R, Fernie AR. 2016. The regulation of essential amino acid synthesis and accumulation in plants. Annual review of plant biology 67: 153–178.

Gemmer MR, Richter C, Schmutzer T, Raorane ML, Junker B, Pillen K, Maurer A. 2021. Genome-wide association study on metabolite accumulation in a wild barley NAM population reveals natural variation in sugar metabolism. Plos one 16: e0246510.

Girard AL, Awika JM. 2018. Sorghum polyphenols and other bioactive components as functional and health promoting food ingredients. Journal of Cereal Science 84: 112–124.

Gong L, Chen W, Gao Y, Liu X, Zhang H, Xu C, Yu S, Zhang Q, Luo J. 2013. Genetic analysis of the metabolome exemplified using a rice population. Proceedings of the National Academy of Sciences 110: 20320–20325.

Goodstein DM, Shu S, Howson R, Neupane R, Hayes RD, Fazo J, Mitros T, Dirks W, Hellsten U, Putnam N. 2012. Phytozome: a comparative platform for green plant genomics. Nucleic acids research 40: D1178–D1186.

Gorelick R. 2006. Combining richness and abundance into a single diversity index using matrix analogues of Shannon’s and Simpson’s indices. Ecography 29: 525–530.

Goufo P, Trindade H. 2014. Rice antioxidants: phenolic acids, flavonoids, anthocyanins, proanthocyanidins, tocopherols, tocotrienols, γcoryzanol, and phytic acid. Food science & nutrition 2: 75–104.

Grace SC. 2005. Phenolics as antioxidants. Antioxidants and reactive oxygen species in plants: 141–168.

Habyarimana E, Dall’Agata M, De Franceschi P, Baloch FS. 2019. Genome-wide association mapping of total antioxidant capacity, phenols, tannins, and flavonoids in a panel of Sorghum bicolor and S. bicolor× S. halepense populations using multi-locus models. PLoS One 14: e0225979.

Hajian-Tilaki K. 2013. Receiver operating characteristic (ROC) curve analysis for medical diagnostic test evaluation. Caspian journal of internal medicine 4: 627.

Halliday FW, Rohr JR. 2019. Measuring the shape of the biodiversity-disease relationship across systems reveals new findings and key gaps. Nature communications 10: 5032.

Hamade K, Fliniaux O, Fontaine J-X, Molinié R, Petit L, Mathiron D, Sarazin V, Mesnard F. 2024. NMR and LC–MS-based metabolomics to investigate the efficacy of a commercial bio stimulant for the treatment of wheat (Triticum aestivum). Metabolomics 20: 58.

Hamblin MT, Salas Fernandez MG, Tuinstra MR, Rooney WL, Kresovich S. 2007. Sequence variation at candidate loci in the starch metabolism pathway in sorghum: prospects for linkage disequilibrium mapping. Crop Science 47: S-125-S-134.

Horai H, Arita M, Kanaya S, Nihei Y, Ikeda T, Suwa K, Ojima Y, Tanaka K, Tanaka S, Aoshima K. 2010. MassBank: a public repository for sharing mass spectral data for life sciences. Journal of mass spectrometry 45: 703–714.

Jander G, Joshi V. 2009. Aspartate-derived amino acid biosynthesis in Arabidopsis thaliana. The arabidopsis book/American society of plant biologists 7: e0121.

Kai K, Shimizu B-i, Mizutani M, Watanabe K, Sakata K. 2006. Accumulation of coumarins in Arabidopsis thaliana. Phytochemistry 67: 379–386.

Kanehisa M, Goto S. 2000. KEGG: kyoto encyclopedia of genes and genomes. Nucleic acids research 28: 27–30.

Karakas E, Bulut M, Fernie A. 2025. Metabolome guided treasure hunt-learning from metabolic diversity. Journal of Plant Physiology: 154494.

Kassambara A. 2013. ggplot2: guide to create beautiful graphics in R. Alboukadel KASSAMBARA.

Kathuria D, Thakur S, Singh N. 2024. Advances of metabolomic in exploring phenolic compounds diversity in cereal and their health implications. International Journal of Food Science and Technology 59: 4213–4233.

Kaufman RC, Herald TJ, Bean SR, Wilson JD, Tuinstra MR. 2013. Variability in tannin content, chemistry and activity in a diverse group of tannin containing sorghum cultivars. Journal of the Science of Food and Agriculture 93: 1233–1241.

Kaur S, Kumar K, Singh L, Sharanagat VS, Nema PK, Mishra V, Bhushan B. 2024. Gluten-free grains: Importance, processing and its effect on quality of gluten-free products. Critical reviews in food science and nutrition 64: 1988–2015.

Kautsar SA, Suarez Duran HG, Blin K, Osbourn A, Medema MH. 2017. plantiSMASH: automated identification, annotation and expression analysis of plant biosynthetic gene clusters. Nucleic acids research 45: W55–W63.

Khan A, Tian R, Bean SR, Yerka M, Jiao Y. 2024. Transcriptome and metabolome analyses reveal regulatory networks associated with nutrition synthesis in sorghum seeds. Communications Biology 7: 841.

Kim S. 2015. ppcor: an R package for a fast calculation to semi-partial correlation coefficients. Communications for statistical applications and methods 22: 665.

KobuecLekalake RI, Taylor JR, De Kock HL. 2007. Effects of phenolics in sorghum grain on its bitterness, astringency and other sensory properties. Journal of the Science of Food and Agriculture 87: 1940–1948.

Lam PY, Zhu F-Y, Chan WL, Liu H, Lo C. 2014. Cytochrome P450 93G1 is a flavone synthase II that channels flavanones to the biosynthesis of tricin O-linked conjugates in rice. Plant Physiology 165: 1315–1327.

Lee J, Hwang Y-S, Kim ST, Yoon W-B, Han WY, Kang I-K, Choung M-G. 2017. Seed coat color and seed weight contribute differential responses of targeted metabolites in soybean seeds. Food Chemistry 214: 248–258.

Lee S, Miropolsky L, Wu M, Lee M, CompQuadForm D. 2011. Package “SKAT.”. In CRAN.

Li J, Tang W, Zhang Y-W, Chen K-N, Wang C, Liu Y, Zhan Q, Wang C, Wang S-B, Xie S-Q. 2018. Genome-wide association studies for five forage quality-related traits in sorghum (Sorghum bicolor L.). Frontiers in plant science 9: 1146.

Li M-X, Yeung JM, Cherny SS, Sham PC. 2012. Evaluating the effective numbers of independent tests and significant p-value thresholds in commercial genotyping arrays and public imputation reference datasets. Human genetics 131: 747–756.

Li Q, Fu C, Liang C, Ni X, Zhao X, Chen M, Ou L. 2022. Crop lodging and the roles of lignin, cellulose, and hemicellulose in lodging resistance. Agronomy 12: 1795.

Li Y, Miao Y, Yuan H, Huang F, Sun M, He L, Liu X, Luo J. 2024. Volatilome-based GWAS identifies OsWRKY19 and OsNAC021 as key regulators of rice aroma. Molecular Plant 17: 1866–1882.

Liang J, Shen Q, Wang L, Liu J, Fu J, Zhao L, Xu M, Peters RJ, Wang Q. 2021. Rice contains a biosynthetic gene cluster associated with production of the casbanectype diterpenoid phytoalexin entc10coxodepressin. New Phytologist 231: 85–93.

Liaqat W, Altaf MT, Barutçular C, Mohamed HI, Ahmad H, Jan MF, Khan EH. 2024. Sorghum: a star crop to combat abiotic stresses, food insecurity, and hunger under a changing climate: a review. Journal of Soil Science and Plant Nutrition 24: 74-101.

Liaw A, Wiener M. 2002. Classification and regression by randomForest. R news 2: 18–22.

Lipka AE, Gore MA, Magallanes-Lundback M, Mesberg A, Lin H, Tiede T, Chen C, Buell CR, Buckler ES, Rocheford T. 2013. Genome-wide association study and pathway-level analysis of tocochromanol levels in maize grain. G3: Genes, Genomes, Genetics 3: 1287-1299.

Lipka AE, Tian F, Wang Q, Peiffer J, Li M, Bradbury PJ, Gore MA, Buckler ES, Zhang Z. 2012. GAPIT: genome association and prediction integrated tool. Bioinformatics 28: 2397–2399.

Lippert C, Listgarten J, Liu Y, Kadie CM, Davidson RI, Heckerman D. 2011. FaST linear mixed models for genome-wide association studies. Nature methods 8: 833–835.

Liu X, Huang M, Fan B, Buckler ES, Zhang Z. 2016. Iterative usage of fixed and random effect models for powerful and efficient genome-wide association studies. PLoS genetics 12: e1005767.

López-González C, Juárez-Colunga S, Morales-Elías NC, Tiessen A. 2019. Exploring regulatory networks in plants: transcription factors of starch metabolism. PeerJ 7: e6841.

Luo J. 2015. Metabolite-based genome-wide association studies in plants. Current opinion in plant biology 24: 31–38.

Matros A, Houston K, Tucker MR, Schreiber M, Berger B, Aubert MK, Wilkinson LG, Witzel K, Waugh R, Seiffert U. 2021. Genome-wide association study reveals the genetic complexity of fructan accumulation patterns in barley grain. Journal of Experimental Botany 72: 2383–2402.

Mazumder S, Bhattacharya D, Lahiri D, Moovendhan M, Sarkar T, Nag M. 2024. Harnessing the nutritional profile and health benefits of millets: a solution to global food security problems. Critical Reviews in Food Science and Nutrition: 1–22.

Mbanjo EGN, Kretzschmar T, Jones H, Ereful N, Blanchard C, Boyd LA, Sreenivasulu N. 2020. The genetic basis and nutritional benefits of pigmented rice grain. Frontiers in genetics 11: 229.

Mbanjo EGN, Pasion EA, Jones H, Carandang S, Misra G, Ignacio JC, Kretzschmar T, Sreenivasulu N, Boyd LA. 2023. Unravelling marker trait associations linking nutritional value with pigmentation in rice seed. The plant genome 16: e20360.

McCarthy MI, Abecasis GR, Cardon LR, Goldstein DB, Little J, Ioannidis JP, Hirschhorn JN. 2008. Genome-wide association studies for complex traits: consensus, uncertainty and challenges. Nature reviews genetics 9: 356–369.

McLaren W, Gil L, Hunt SE, Riat HS, Ritchie GR, Thormann A, Flicek P, Cunningham F. 2016. The ensembl variant effect predictor. Genome biology 17: 1–14.

Melville J. 2019. uwot: The uniform manifold approximation and projection (UMAP) method for dimensionality reduction. CRAN: Contributed Packages.

Menni C, Zhu J, Le Roy CI, Mompeo O, Young K, Rebholz CM, Selvin E, North KE, Mohney RP, Bell JT. 2020. Serum metabolites reflecting gut microbiome alpha diversity predict type 2 diabetes. Gut microbes 11: 1632–1642.

Merrick LF, Burke AB, Zhang Z, Carter AH. 2022. Comparison of single-trait and multi-trait genome-wide association models and inclusion of correlated traits in the dissection of the genetic architecture of a complex trait in a breeding program. Frontiers in Plant Science 12: 772907.

Michalski A, Damoc E, Hauschild J-P, Lange O, Wieghaus A, Makarov A, Nagaraj N, Cox J, Mann M, Horning S. 2011. Mass spectrometry-based proteomics using Q Exactive, a high-performance benchtop quadrupole Orbitrap mass spectrometer. Molecular & cellular proteomics 10.

Mokra D, Joskova M, Mokry J. 2022. Therapeutic effects of green tea polyphenol (c)-Epigallocatechin-3-Gallate (EGCG) in relation to molecular pathways controlling inflammation, oxidative stress, and apoptosis. International journal of molecular sciences 24: 340.

Mukherjee A, Maheshwari U, Sharma V, Sharma A, Kumar S. 2024. Functional insight into multi-omics-based interventions for climatic resilience in sorghum (Sorghum bicolor): a nutritionally rich cereal crop. Planta 259: 91.

Nip W, Burns E. 1969. Pigment characterization in grain sorghum. I. Red varieties. Cereal Chem 46: 490–495.

Noel JP, Austin MB, Bomati EK. 2005. Structure–function relationships in plant phenylpropanoid biosynthesis. Current opinion in plant biology 8: 249–253.

Nützmann HW, Huang A, Osbourn A. 2016. Plant metabolic clusters–from genetics to genomics. New phytologist 211: 771–789.

Ojwang LO, Dykes L, Awika JM. 2012. Ultra performance liquid chromatography–tandem quadrupole mass spectrometry profiling of anthocyanins and flavonols in cowpea (Vigna unguiculata) of varying genotypes. Journal of agricultural and food chemistry 60: 3735–3744.

Pallister T, Jackson MA, Martin TC, Zierer J, Jennings A, Mohney RP, MacGregor A, Steves CJ, Cassidy A, Spector TD. 2017. Hippurate as a metabolomic marker of gut microbiome diversity: Modulation by diet and relationship to metabolic syndrome. Scientific reports 7: 13670.

Park JS, Rho HS, Kim DH, Chang IS. 2006. Enzymatic preparation of kaempferol from green tea seed and its antioxidant activity. Journal of agricultural and food chemistry 54: 2951–2956.

Paterson AH, Bowers JE, Bruggmann R, Dubchak I, Grimwood J, Gundlach H, Haberer G, Hellsten U, Mitros T, Poliakov A. 2009. The Sorghum bicolor genome and the diversification of grasses. Nature 457: 551–556.

Pelletier MK, Shirley BW. 1996. Analysis of flavanone 3-hydroxylase in Arabidopsis seedlings (Coordinate regulation with chalcone synthase and chalcone isomerase). Plant physiology 111: 339–345.

Pence HE, Williams A. 2010. ChemSpider: an online chemical information resource. ACS Publications.

Pontieri P, Pepe G, Campiglia P, Merciai F, Basilicata MG, Smolensky D, Calcagnile M, Troisi J, Romano R, Del Giudice F. 2021. Comparison of content in phenolic compounds and antioxidant capacity in grains of white, red, and black sorghum varieties grown in the mediterranean area. ACS Food Science & Technology 1: 1109–1119.

Przybylska-Balcerek A, Frankowski J, Stuper-Szablewska K. 2019. Bioactive compounds in sorghum. European Food Research and Technology 245: 1075–1080.

Qin Sh, Yan F, E S, Xiong P, Tang Sn, Yu Kq, Zhang M, Cheng Yc, Cai W. 2022. Comprehensive characterization of multiple components of Ziziphus jujuba Mill using UHPLCcQcExactive Orbitrap Mass Spectrometers. Food Science & Nutrition 10: 4270–4295.

Rai M, Dutta M, Saito K, Rai A. 2025. A deep dive into plant metabolomics: Milestones, technologies, and translational impact. Plant Physiology 199: kiaf408.

Rao S, Santhakumar AB, Chinkwo KA, Wu G, Johnson SK, Blanchard CL. 2018. Characterization of phenolic compounds and antioxidant activity in sorghum grains. Journal of Cereal Science 84: 103–111.

Ren Z, Fang M, Muhae-Ud-Din G, Gao H, Yang Y, Liu T, Chen W, Gao L. 2021. Metabolomics analysis of grains of wheat infected and noninfected with Tilletia controversa Kühn. Scientific Reports 11: 18876.

Rhodes DH, Hoffmann Jr L, Rooney WL, Herald TJ, Bean S, Boyles R, Brenton ZW, Kresovich S. 2017. Genetic architecture of kernel composition in global sorghum germplasm. BMC genomics 18: 15.

Ribbenstedt A, Ziarrusta H, Benskin JP. 2018. Development, characterization and comparisons of targeted and non-targeted metabolomics methods. PloS one 13: e0207082.

Salgado AL, Glassmire AE, Sedio BE, Diaz R, Stout MJ, Čuda J, Pyšek P, Meyerson LA, Cronin JT. 2023. Metabolomic evenness underlies intraspecific differences among lineages of a wetland grass. Journal of Chemical Ecology 49: 437–450.

Sansone S-A, Fan T, Goodacre R, Griffin JL, Hardy NW, Kaddurah-Daouk R, Kristal BS, Lindon J, Mendes P, Morrison N et al. 2007. The Metabolomics Standards Initiative. Nature Biotechnology 25: 846–848.

Sauvage C, Segura V, Bauchet G, Stevens R, Do PT, Nikoloski Z, Fernie AR, Causse M. 2014. Genome-wide association in tomato reveals 44 candidate loci for fruit metabolic traits. Plant physiology 165: 1120–1132.

Sawada Y, Akiyama K, Sakata A, Kuwahara A, Otsuki H, Sakurai T, Saito K, Hirai MY. 2009. Widely targeted metabolomics based on large-scale MS/MS data for elucidating metabolite accumulation patterns in plants. Plant and Cell Physiology 50: 37–47.

Shannon P, Markiel A, Ozier O, Baliga NS, Wang JT, Ramage D, Amin N, Schwikowski B, Ideker T. 2003. Cytoscape: a software environment for integrated models of biomolecular interaction networks. Genome research 13: 2498.

Shurubor YI, Matson WR, Martin RJ, Kristal BS. 2005. Relative contribution of specific sources of systematic errors and analytical imprecision to metabolite analysis by HPLC–ECD. Metabolomics 1: 159–168.

Shurubor YI, Matson WR, Willett WC, Hankinson SE, Kristal BS. 2007. Biological variability dominates and influences analytical variance in HPLC-ECD studies of the human plasma metabolome. BMC Clinical Pathology 7: 9.

Siddique I. 2021. Unveiling the power of high-performance liquid chromatography: Techniques, applications, and innovations. European Journal of Advances in Engineering and Technology 8: 79–84.

Simeone MLF, Parrella RA, Schaffert RE, Damasceno CM, Leal MC, Pasquini C. 2017. Near infrared spectroscopy determination of sucrose, glucose and fructose in sweet sorghum juice. Microchemical Journal 134: 125–130.

Sindelar M, Patti GJ. 2020. Chemical discovery in the era of metabolomics. Journal of the American Chemical Society 142: 9097–9105.

Smith CA, Want EJ, O’Maille G, Abagyan R, Siuzdak G. 2006. XCMS: processing mass spectrometry data for metabolite profiling using nonlinear peak alignment, matching, and identification. Analytical chemistry 78: 779–787.

Song S, Zhang L, Zhao Y, Sheng C, Zhou W, Dossou SSK, Wang L, You J, Zhou R, Wei X. 2022. Metabolome genomecwide association study provides biochemical and genetic insights into natural variation of primary metabolites in sesame. The Plant Journal 112: 1051–1069.

Spicer RA, Salek R, Steinbeck C. 2017. A decade after the metabolomics standards initiative it’s time for a revision. Scientific Data 4: 1–3.

Su XiaoYu SX, Rhodes D, Xu JingWen XJ, Chen Xi CX, Davis H, Wang DongHai WD, Herald TJ, Wang WeiQun WW. 2017. Phenotypic diversity of anthocyanins in sorghum accessions with various pericarp pigments.

Sytar O, Zivcak M, Neugart S, Toutounchi PM, Brestic M. 2019. Precultivation of young seedlings under different color shades modifies the accumulation of phenolic compounds in Cichorium leaves in later growth phases. Environmental and Experimental Botany 165: 30-38.

Tam V, Patel N, Turcotte M, Bossé Y, Paré G, Meyre D. 2019. Benefits and limitations of genome-wide association studies. Nature Reviews Genetics 20: 467–484.

Tanwar R, Panghal A, Chaudhary G, Kumari A, Chhikara N. 2023. Nutritional, phytochemical and functional potential of sorghum: A review. Food Chemistry Advances 3: 100501.

Team G, Mesnard T, Hardin C, Dadashi R, Bhupatiraju S, Pathak S, Sifre L, Rivière M, Kale MS, Love J. 2024. Gemma: Open models based on gemini research and technology. arXiv preprint arXiv:240308295.

Thakur NR, Gorthy S, Vemula A, Odeny DA, Ruperao P, Sargar PR, Mehtre SP, Kalpande HV, Habyarimana E. 2024. Genome-wide association study and expression of candidate genes for Fe and Zn concentration in sorghum grains. Scientific Reports 14: 12729.

Thitisaksakul M, Jiménez RC, Arias MC, Beckles DM. 2012. Effects of environmental factors on cereal starch biosynthesis and composition. Journal of cereal science 56: 67–80.

Tibbs Cortes L, Zhang Z, Yu J. 2021. Status and prospects of genomecwide association studies in plants. The plant genome 14: e20077.

Tiozon RJN, Sreenivasulu N, Alseekh S, Sartagoda KJD, Usadel B, Fernie AR. 2023. Metabolomics and machine learning technique revealed that germination enhances the multi-nutritional properties of pigmented rice. Communications biology 6: 1000.

Upadhyaya HD, Sharma S, Dwivedi SL, Singh SK. 2014. Sorghum genetic resources: conservation and diversity assessment for enhanced utilization in sorghum improvement.

Vanamala JK, Massey AR, Pinnamaneni SR, Reddivari L, Reardon KF. 2018. Grain and sweet sorghum (Sorghum bicolor L. Moench) serves as a novel source of bioactive compounds for human health. Critical Reviews in Food Science and Nutrition 58: 2867–2881.

Volk GM, Richards CM. 2008. Availability of genotypic data for USDA-ARS National Plant Germplasm System accessions using the genetic resources information network (GRIN) database. HortScience 43: 1365–1366.

Von Mering C, Jensen LJ, Snel B, Hooper SD, Krupp M, Foglierini M, Jouffre N, Huynen MA, Bork P. 2005. STRING: known and predicted protein–protein associations, integrated and transferred across organisms. Nucleic acids research 33: D433–D437.

Voorman A, Lumley T, McKnight B, Rice K. 2011. Behavior of QQ-plots and genomic control in studies of gene-environment interaction. PloS one 6: e19416.

Wang Y, Wang X, Sun S, Jin C, Su J, Wei J, Luo X, Wen J, Wei T, Sahu SK. 2022. GWAS, MWAS and mGWAS provide insights into precision agriculture based on genotype-dependent microbial effects in foxtail millet. Nature Communications 13: 5913.

Wen W, Li D, Li X, Gao Y, Li W, Li H, Liu J, Liu H, Chen W, Luo J. 2014. Metabolome-based genome-wide association study of maize kernel leads to novel biochemical insights. Nature communications 5: 3438.

Wickham H. 2011. ggplot2. Wiley interdisciplinary reviews: computational statistics 3: 180–185.

Wishart DS, Guo A, Oler E, Wang F, Anjum A, Peters H, Dizon R, Sayeeda Z, Tian S, Lee BL. 2022. HMDB 5.0: the human metabolome database for 2022. Nucleic acids research 50: D622–D631.

Wu G, Johnson SK, Bornman JF, Bennett SJ, Clarke MW, Singh V, Fang Z. 2016. Growth temperature and genotype both play important roles in sorghum grain phenolic composition. Scientific reports 6: 21835.

Wu MC, Lee S, Cai T, Li Y, Boehnke M, Lin X. 2011. Rare-variant association testing for sequencing data with the sequence kernel association test. The American Journal of Human Genetics 89: 82–93.

Wu Y, Guo T, Mu Q, Wang J, Li X, Wu Y, Tian B, Wang ML, Bai G, Perumal R. 2019. Allelochemicals targeted to balance competing selections in African agroecosystems. Nature plants 5: 1229–1236.

Wu Y, Li X, Xiang W, Zhu C, Lin Z, Wu Y, Li J, Pandravada S, Ridder DD, Bai G. 2012. Presence of tannins in sorghum grains is conditioned by different natural alleles of Tannin1. Proceedings of the National Academy of Sciences 109: 10281–10286.

Wurtzel ET, Kutchan TM. 2016. Plant metabolism, the diverse chemistry set of the future. Science 353: 1232–1236.

Xiao R, Ma Y, Zhang D, Qian L. 2018. Discrimination of conventional and organic rice using untargeted LC-MS-based metabolomics. Journal of Cereal Science 82: 73–81.

Xu Z, Dai J, Kang T, Shah K, Li Q, Liu K, Xing L, Ma J, Zhang D, Zhao C. 2022. PpePL1 and PpePL15 are the core members of the pectate lyase gene family involved in peach fruit ripening and softening. Frontiers in Plant Science 13: 844055.

Yin L, Zhang H, Tang Z, Xu J, Yin D, Zhang Z, Yuan X, Zhu M, Zhao S, Li X. 2021. rMVP: a memory-efficient, visualization-enhanced, and parallel-accelerated tool for genome-wide association study. Genomics, Proteomics and Bioinformatics 19: 619–628.

Young AS. 2024. Genome-wide association studies have problems due to confounding: Are family-based designs the answer? PLoS biology 22: e3002568.

Zhang R, Jia G, Diao X. 2023a. geneHapR: an R package for gene haplotypic statistics and visualization. BMC bioinformatics 24: 199.

Zhang T, Lu L, Yang N, Fisk ID, Wei W, Wang L, Li J, Sun Q, Zeng R. 2023b. Integration of hyperspectral imaging, non-targeted metabolomics and machine learning for vigour prediction of naturally and accelerated aged sweetcorn seeds. Food Control 153: 109930.

Zhang X, Tian X, Gao X, Sun G, Peng Z, Geng X, Peng J, Dai P, Wang X, Li H. 2025. Integrated metabolomic and transcriptomic analyses identify MYB genes regulating key metabolites and agronomic traits in upland cotton Gossypium hirsutum. Nature Genetics: 1–12.

Zhang Y, Li M, Gao H, Wang B, Tongcheng X, Gao B, Yu L. 2019. Triacylglycerol, fatty acid, and phytochemical profiles in a new red sorghum variety (Ji Liang No. 1) and its antioxidant and anticinflammatory properties. Food Science & Nutrition 7: 949-958.

Zhang Z, Ersoz E, Lai C-Q, Todhunter RJ, Tiwari HK, Gore MA, Bradbury PJ, Yu J, Arnett DK, Ordovas JM. 2010. Mixed linear model approach adapted for genome-wide association studies. Nature genetics 42: 355–360.

Zheng L-Y, Guo X-S, He B, Sun L-J, Peng Y, Dong S-S, Liu T-F, Jiang S, Ramachandran S, Liu C-M. 2011. Genome-wide patterns of genetic variation in sweet and grain sorghum (Sorghum bicolor). Genome biology 12: 1–15.

Zhou S, Kremling KA, Bandillo N, Richter A, Zhang YK, Ahern KR, Artyukhin AB, Hui JX, Younkin GC, Schroeder FC. 2019. Metabolome-scale genome-wide association studies reveal chemical diversity and genetic control of maize specialized metabolites. The Plant Cell 31: 937–955.

Zhou Z, Luo M, Zhang H, Yin Y, Cai Y, Zhu Z-J. 2022. Metabolite annotation from knowns to unknowns through knowledge-guided multi-layer metabolic networking. Nature communications 13: 6656.

Zhu G, Wang S, Huang Z, Zhang S, Liao Q, Zhang C, Lin T, Qin M, Peng M, Yang C. 2018. Rewiring of the fruit metabolome in tomato breeding. Cell 172: 249–261. e212.

Zhu X, Galili G. 2003. Increased lysine synthesis coupled with a knockout of its catabolism synergistically boosts lysine content and also transregulates the metabolism of other amino acids in Arabidopsis seeds. The Plant Cell 15: 845–853.

